# Longitudinal Analysis of Particulate Air Pollutants and Adolescent Delinquent Behavior in Southern California

**DOI:** 10.1101/208793

**Authors:** Diana Younan, Catherine Tuvblad, Meredith Franklin, Fred Lurmann, Lianfa Li, Jun Wu, Kiros Berhane, Laura A. Baker, Jiu-Chiuan Chen

## Abstract

Animal experiments and cross-sectional human studies have linked particulate matter (PM) with increased behavioral problems. We conducted a longitudinal study to examine whether the trajectories of delinquent behavior are affected by PM_2.5_ (PM with aerodynamic diameter ≤2.5 μm) exposures before and during adolescence. We used the parent-reported Child Behavior Checklist at age 9-18 with repeated measures every ~2-3 years (up to 4 behavioral assessments) on 682 children from the Risk Factors for Antisocial Behavior Study conducted in a multi-ethnic cohort of twins born in 1990-1995. Based on prospectively-collected residential addresses and a spatiotemporal model of ambient air concentrations in Southern California, monthly PM_2.5_ estimates were aggregated to represent long-term (1-, 2-, 3-year average) exposures preceding baseline and cumulative average exposure until the last assessment. Multilevel mixed-effects models were used to examine the association between PM_2.5_ exposure and individual trajectories of delinquent behavior, adjusting for within-family/within-individual correlations and potential confounders. We also examined whether psychosocial factors modified this association. The results suggest that PM_2.5_ exposure at baseline and cumulative exposure during follow-up was significantly associated (p<0.05) with increased delinquent behavior. The estimated effect sizes (per interquartile increase of PM_2.5_ by 3.12-5.18 µg/m^3^) were equivalent to the difference in delinquency scores between adolescents who are 3.5-4 years apart in age. The adverse effect was stronger in families with unfavorable parent-to-child relationships, increased parental stress or maternal depressive symptoms. Overall, these findings suggest long-term PM_2.5_ exposure may increase delinquent behavior of urban-dwelling adolescents, with the resulting neurotoxic effect aggravated by psychosocial adversities.

---

Delinquency, defined as behaviors and attitudes that violate societal norms, values, and laws, is a strong predictor of future criminal and antisocial activities (Murray and Farrington 2010). Early delinquency predicts negative outcomes later in life, including academic underachievement, unemployment, mental disorders, substance use, and dysfunctional families (Murray and Farrington 2010). As youth incarceration costs state and local governments $8-12 billion annually (Petteruti 2011), delinquent behavior, if not appropriately intervened, may amount to a significant economic and social burden.

Adolescence is an important developmental period characterized by significant social, biological, and physiological changes. This phase of functional and structural change in the prefrontal lobe is a crucial time for shaping behavioral trajectories (Carroll et al. 2014), and is a vulnerable period during which developmental processes may be easily disrupted (Rauh et al. 2010). Delinquent behavior increases during this critical time and peaks during mid-adolescence (Murray and Farrington 2010), therefore, adolescents are an important population to target for early intervention strategies. Approximately 55-87% of the total variance of delinquency is attributable to environment (Burt 2009), but previous studies have mainly focused on social factors and overlooked the influence of physical environments. Rates of juvenile delinquency are highest in urban neighborhoods (Shaw and McKay 1942), highlighting the need to identify modifiable environmental factors in these areas.

Only few environmental neurotoxicants have been implicated as modifiable risk factors for delinquent behavior. Existing literature supports an association between prenatal secondhand smoke and externalizing problems, including delinquency (Tiesler and Heinrich 2014). Early-life exposure to lead increased delinquent behavior in pre-adolescent boys and predicted adjudicated delinquency in adolescents (Needleman et al. 1996). Over the last 15 years, both experimental animal models and epidemiologic studies have reported developmental neurotoxicity of ambient air pollutants (Block et al. 2012), especially particulate matter (PM). Air pollution is the most abundant source of environmentally induced inflammation and oxidative stress, and recent reports have revealed that small particles may enter the brain contributing to structural damage and neurodegeneration (Maher et al. 2016). Cross-sectional studies (Forns et al. 2015; Haynes et al. 2011; Perera et al. 2013; Peterson et al. 2015; Yorifuji et al. 2016) have examined the association of behavioral problems with particulate air pollutants. Although PM_2.5_ (PM with aerodynamic diameter < 2.5-µm) exposure is common in urban areas, very little is known about its influence on delinquent behavior, especially over time. This longitudinal study aims to investigate whether the individual trajectories of delinquent behavior are affected by residential exposure to ambient air pollutants in urban-dwelling adolescents from Southern California, a region where outdoor PM_2.5_ concentrations often exceed state and national annual standards (South Coast Air Quality Management District 2013). Our secondary aim was to explore whether social stressors may enhance a child’s susceptibility to the putative adverse behavioral effects of PM_2.5_, as these possible stress-pollution interactions have become increasing recognized in the literature (Cooney 2011).

## Methods

### Study Design

Participants were drawn from the Risk Factors for Antisocial Behavior (RFAB) twin study (Baker et al. 2007). Families were recruited from Los Angeles and surrounding counties, representative of the multi-ethnic and socioeconomically-diverse population of the greater Los Angeles area (Baker et al. 2007). The initial cohort included over 780 monozygotic and dizygotic (same-sex and opposite sex) twin pairs and triplets born in 1990-1995 and aged 9-10 years at the RFAB inception in 2000. Study protocols were approved by the Institutional Review Board at the University of Southern California (USC). Informed consent was obtained from all individual participants included in the study.

The present study used data collected with up to four behavioral assessments from childhood to adolescence (Baker et al. 2007). Our study base was defined as subjects with at least two assessments of delinquent behavior at age 9-18 (*n* = 1299). For eligibility, children had to provide valid residential addresses at the scheduled testing dates, and their co-twin or triplet siblings also had to participate. Effects of early-life exposure were not studied, because ambient PM_2.5_ data were not collected until 1999. However, comparing the putative effects of long-term exposure estimated over an extended period before behavioral assessments started may provide valuable insights to understanding the contribution of more recent exposure (e.g., 1-year average) versus remote exposure (e.g., 3-year average) affecting the trajectories of delinquent behavior. Therefore, the analytic sample was limited to participants with complete data on PM_2.5_ exposures (1-, 2-, and 3-year averages) before the baseline assessment. A total of 682 subjects (from 338 families) met these criteria (Figure 1). The restriction of exposure data was not extended beyond 3 years, as it would greatly reduce sample size and statistical power.

**Figure 1.**
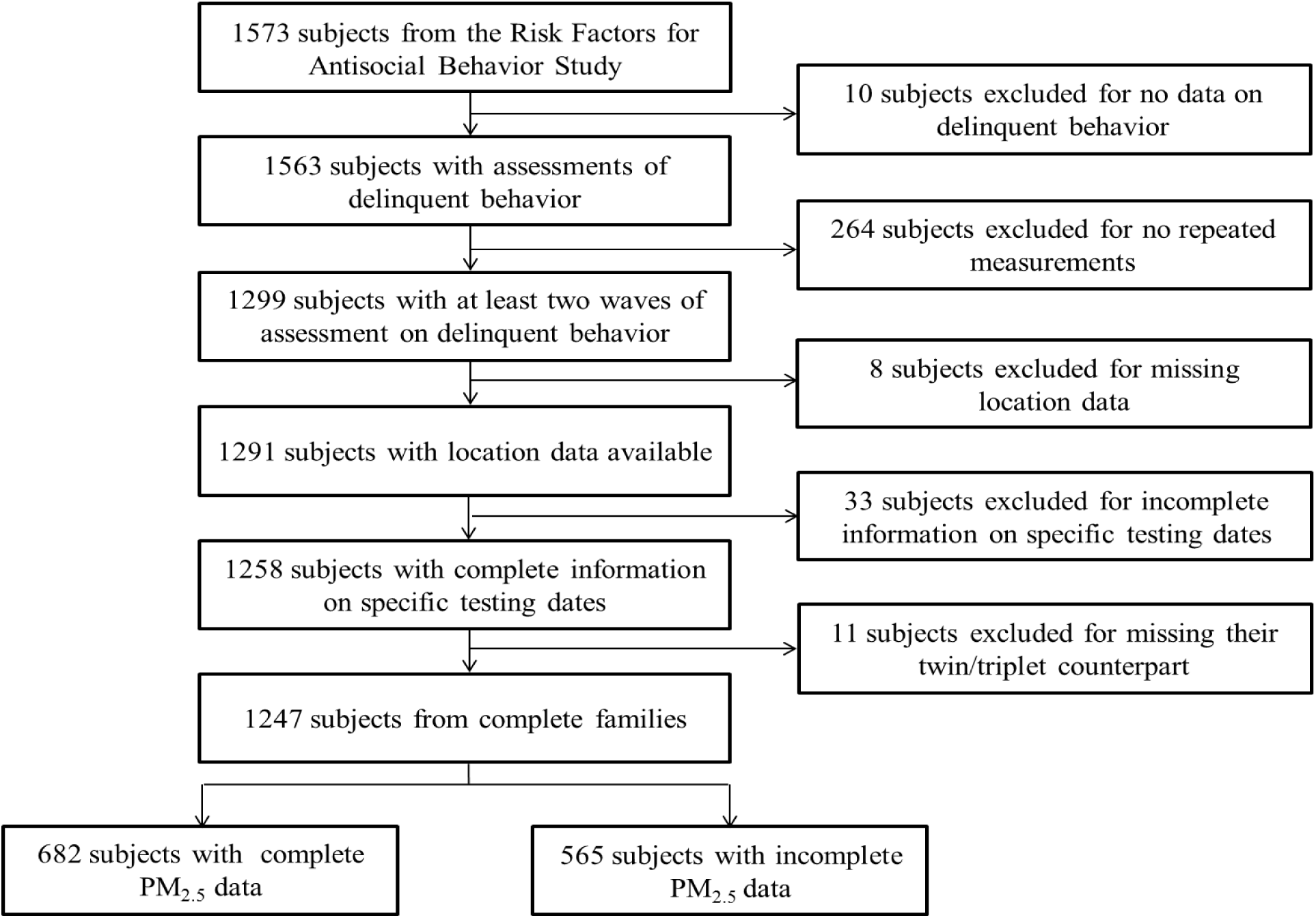
Study flow chart indicating selection of study participants for current analyses.

### Behavioral Assessment

Delinquent behavior over the preceding 6 months was assessed with the parent-reported version of the widely used *Child Behavior Checklist (CBCL/6-18)*. The high reliability and validity of the CBCL has been reported elsewhere (Achenbach and Rescorla 2001). The *Delinquent (Rule-Breaking) Behavior* subscale consists of 13 items, including lying and cheating, feeling no guilt, truancy, stealing, vandalism, running away, fire setting, and substance use. Each item was scored on a 3-point scale (0: not true; 1: sometimes true; and 2: very true/often true), and a continuous raw score was created by summing across items. The CBCL assessment had a relatively high internal consistency (average Cronbach’s Alpha: α = 0.71) in the RFAB cohort across 4 assessments (mean age ± SD: 9.22 ± 0.42; 11.32 ± 0.92; 14.49 ± 0.76; 16.70 ± 0.66).

### Estimation of Ambient Air Pollution Exposure

#### Residential location data and geocoding

Residential addresses, prospectively collected through parent-reports at each wave, were sent to the USC Spatial Sciences Institute for geocoding, which followed standard procedures and returned high-quality data, with successful matching by exact parcel locations or specific street segments for 98.6% of RFAB families (see baseline geographic distribution presented in Figure S1). The remaining addresses were geocoded satisfactorily with Google Earth based on visual acceptance.

#### Spatiotemporal generalized additive modeling for PM_2.5_

Daily PM_2.5_ concentrations recorded at twenty-five monitors in our study area were acquired in 2000-2014. Cross-validated spatiotemporal generalized additive models (Wood 2006) were fit to monthly average concentrations in overlapping five-year (i.e., 2000-2005, 2004-2008, 2007-2011, 2010-2014) segments at each monitor (see Supplement for more details). These models (with an average model cross-validation R^2^=0.71) were then applied to estimating PM_2.5_ concentrations. From the resulting monthly exposure time-series in 2000-2014 at each geocoded home location, corresponding exposure estimates were aggregated to represent the average PM_2.5_ 1-, 2-, and 3-years preceding baseline (i.e., the first valid CBCL assessment), as well as the cumulative exposure over follow-up.

#### CALINE freeway NO_x_

We estimated near-roadway exposure to nitrogen oxides (NO_x_), using the CALINE4 dispersion model (Benson 1992), which incorporates roadway geometry, traffic volume and emission rate by roadway link, and meteorological conditions. Average ambient NO_x_ concentrations from local (within 5-km) traffic were obtained during the testing year, and cumulative exposures over follow-up were also assigned (see Supplement).

#### Relevant Covariate Data

A directed acyclic graph (DAG) (Howards et al. 2012) was used to identify potential confounders (Figure S2) known to predict delinquency and likely influence where people lived (and thus their exposure to ambient air pollution), including age, gender, race/ethnicity, household socioeconomic status (SES), neighborhood socioeconomic (nSES) characteristics (defined by US census data), and self-perceived neighborhood quality. Potential confounding by other covariates, including freeway NO_x_, traffic density (proxy for traffic noise), neighborhood greenspace, meteorological factors (relative humidity; temperature), urbanicity, prenatal secondhand smoke and maternal depressive symptoms (proxies for early-life and maternal risk factors) were also evaluated by DAG because they correlated with PM_2.5_ or increased delinquent behavior. Also included were measures of parent-to-child affect (PCA), perceived parental stress and maternal depressive symptoms (by standardized/validated instruments; see Supplement) as potential moderators.

#### Statistical Analysis

Three-level mixed effects models (Diggle et al. 2002) with the restricted maximum likelihood and an unstructured covariance structure were constructed by regressing continuous raw delinquency scores on air pollution exposures (1-, 2-, and 3-years prior to baseline; cumulative exposure over follow-up), while accounting for within-family (random intercept and slope [age]) and within-individual (random intercept) correlations and multiple potential confounders, including age as a time-varying covariate (see Supplement). Separate analyses were conducted to investigate the independent effects of air pollution at each exposure temporal scale. Further analyses were also conducted to investigate whether air pollution exposures might affect the slope of delinquent behavior over time (the age slope). We ran sensitivity analyses to evaluate other potential confounding and conducted exploratory analyses on possible effect modification (see Supplement). All analyses were performed using SAS (version 9.4) and figures were created using R software (version 2.15.2).

## Results

Descriptive statistics of our main exposure and outcome of interest are presented in Table S1. Our study sample (*n* = 682) and the excluded subjects (*n* = 617) had similar delinquent behavior scores and did not significantly differ by gender, ethnicity, household SES, nSES, neighborhood quality, or freeway NO_x_ (Table S2). Subjects included in our analyses were slightly older (p < 0.0001), with more entering the cohort at Wave 3 (2006-2010) when PM_2.5_ exposure was lower (p < 0.0001).

Subjects who were enrolled earlier, girls, racial/ethnic minority groups (e.g., non-Hispanic whites; African American), from households of lower SES, living in urban areas, neighborhoods with unfavorable nSES characteristics or perceiving poorer neighborhood quality, had higher levels of ambient PM_2.5_ exposures, as compared to their counterparts (Table 1). Higher PM_2.5_ estimates were also observed in locations with elevated NO_x_ from freeway, limited greenspace, and higher temperatures and relative humidity. Parents with higher PM_2.5_ exposures perceived the highest level of stress, but tended to have more favorable PCA, except for the subscale of parent-reported negative PCA. Mothers of twins with higher PM_2.5_ exposures reported more depressive symptoms.

**Table 1.** Population Characteristics at Baseline in Relation to PM_2.5_ Exposure 1-year Prior to Baseline

| Population Characteristics<sup>b</sup> | Quartile of PM<sub>2.5</sub> (ug/m<sup>3</sup>) |  |  |  |  |
| --- | --- | --- | --- | --- | --- |
|  | 7.3-14.9<br>Median = 13.2<br>(n = 171) | 14.9-18.3<br>Median = 16.1<br>(n = 170) | 18.3-20.5<br>Median = 19.4<br>(n = 170) | 20.5-23.2<br>Median = 20.9<br>(n = 171) |  |
| **Categorical variables** | **n (%)** | **n (%)** | **n (%)** | **n (%)** | **p-value<sup>c</sup>** |
| Gender |  |  |  |  | <0.01 |
| Boys | 101 (31.0) | 79 (24.2) | 76 (23.3) | 70 (21.5) |  |
| Girls | 70 (19.7) | 91 (25.6) | 94 (26.4) | 101 (28.4) |  |
| Ethnicity |  |  |  |  | <0.01 |
| White | 84 (37.3) | 67 (29.8) | 49 (21.8) | 25 (11.1) |  |
| Hispanic | 32 (14.8) | 40 (18.5) | 52 (24.1) | 92 (42.6) |  |
| African American | 10 (11.6) | 18 (20.9) | 28 (32.6) | 30 (34.9) |  |
| Mixed | 41 (32.8) | 31 (24.8) | 31 (24.8) | 22 (17.6) |  |
| Other or missing | 4 (13.3) | 14 (46.7) | 10 (33.3) | 2 (6.7) |  |
| Prenatal SHS exposure |  |  |  |  | <0.01 |
| No | 147 (27.8) | 136 (25.7) | 119 (22.5) | 127 (24.0) |  |
| Yes | 16 (13.8) | 26 (22.4) | 40 (34.5) | 34 (29.3) |  |
| Urbanicity |  |  |  |  | <0.01 |
| Non-urban areas | 10 (100%) | 0 (0%) | 0 (0%) | 0 (0%) |  |
| Urban areas | 161 (24.0%) | 170 (25.3%) | 170 (25.3%) | 171 (25.5%) |  |
| **Continuous variables** | **Mean (SD)** | **Mean (SD)** | **Mean (SD)** | **Mean (SD)** | **p-value<sup>d</sup>** |
| Age, y | 13.9 (1.9) | 12.1 (2.4) | 9.5 (0.7) | 9.6 (0.6) | <0.01 |
| Household SES | 43.2 (8.7) | 44.6 (11.3) | 43.2 (13.3) | 39.3 (12.6) | <0.01 |
| Neighborhood SES | 0.4 (0.9) | 0.3 (1.0) | 0.1 (0.9) | -0.5 (0.7) | <0.01 |
| Neighborhood quality | 51.4 (8.9) | 51.1 (8.4) | 50.5 (9.4) | 45.1 (13.5) | <0.01 |
| Freeway NO<sub>x</sub>, ppb | 11.1 (15.0) | 16.9 (21.6) | 21.4 (18.3) | 23.6 (16.0) | <0.01 |
| Traffic density | 73.8 (153.5) | 73.7 (124.7) | 82.5 (124.7) | 88.4 (146.2) | 0.69 |
| Neighborhood greenspace | 0.3 (0.1) | 0.3 (0.1) | 0.3 (0.1) | 0.3 (0.1) | <0.01 |
| Temperature, °C | 17.5 (0.9) | 17.5 (0.8) | 17.4 (0.7) | 17.9 (0.6) | <0.01 |
| Relative humidity, % | 57.0 (9.3) | 59.7 (8.1) | 66.6 (7.1) | 64.3 (4.9) | <0.01 |
| Self-reported (-) PCA | 2.5 (0.7) | 2.4 (0.7) | 2.2 (0.8) | 2.2 (0.7) | <0.01 |
| Self-reported (+) PCA | 3.6 (0.6) | 3.7 (0.6) | 3.9 (0.6) | 3.9 (0.6) | <0.01 |
| Parent-reported (-) PCA | 2.3 (0.6) | 2.4 (0.6) | 2.3 (0.6) | 2.3 (0.5) | 0.22 |
| Parent-reported (+) PCA | 4.0 (0.4) | 4.0 (0.4) | 4.2 (0.4) | 4.2 (0.4) | <0.01 |
| Parental stress | 30.0 (8.0) | 29.7 (8.3) | 32.6 (9.9) | 34.2 (8.3) | <0.01 |
| Maternal depression | 0.19 (0.33) | 0.18 (0.29) | 0.35 (0.54) | 0.37 (0.48) | <0.01 |

Abbreviations: NO<sub>x</sub>, nitrogen oxides; PCA, parent-to-child affect; PM<sub>2.5</sub>, particulate matter with aerodynamic diameter < 2.5 µm; SES, socioeconomic status; SHS, secondhand smoke.
<sup>a</sup>Total number of subjects decreases slightly due to missing values.
<sup>b</sup>SES: high scores corresponding to higher SES levels; neighborhood quality: higher score for a more positive perception of neighborhood quality; neighborhood greenspace: measured by Normalized Difference Vegetation Index, with higher scores representing more vegetation; PCA: higher positive scores represent better relationships and higher negative scores represent worse relationships.
<sup>c</sup>P-value for Pearson $\chi^2$ test comparing the distribution of PM<sub>2.5</sub> across population characteristics.
<sup>d</sup>P-value for ANOVA test comparing means of population characteristics for quartile of PM<sub>2.5</sub>.

Delinquent behavior increased during adolescence. According to the intra-class correlation, 42% of the variability in delinquency was attributable to between-family differences, with the remaining 58% from within-family differences. More delinquent behavior was found in boys, African Americans, lower SES household, families perceiving poorer neighborhood quality or those living with unfavorable nSES or limited greenspace, as compared to their counterparts (Table 2). Delinquency increased with more unfavorable PCA, higher levels of parental stress, and maternal depressive symptoms.

**Table 2.**
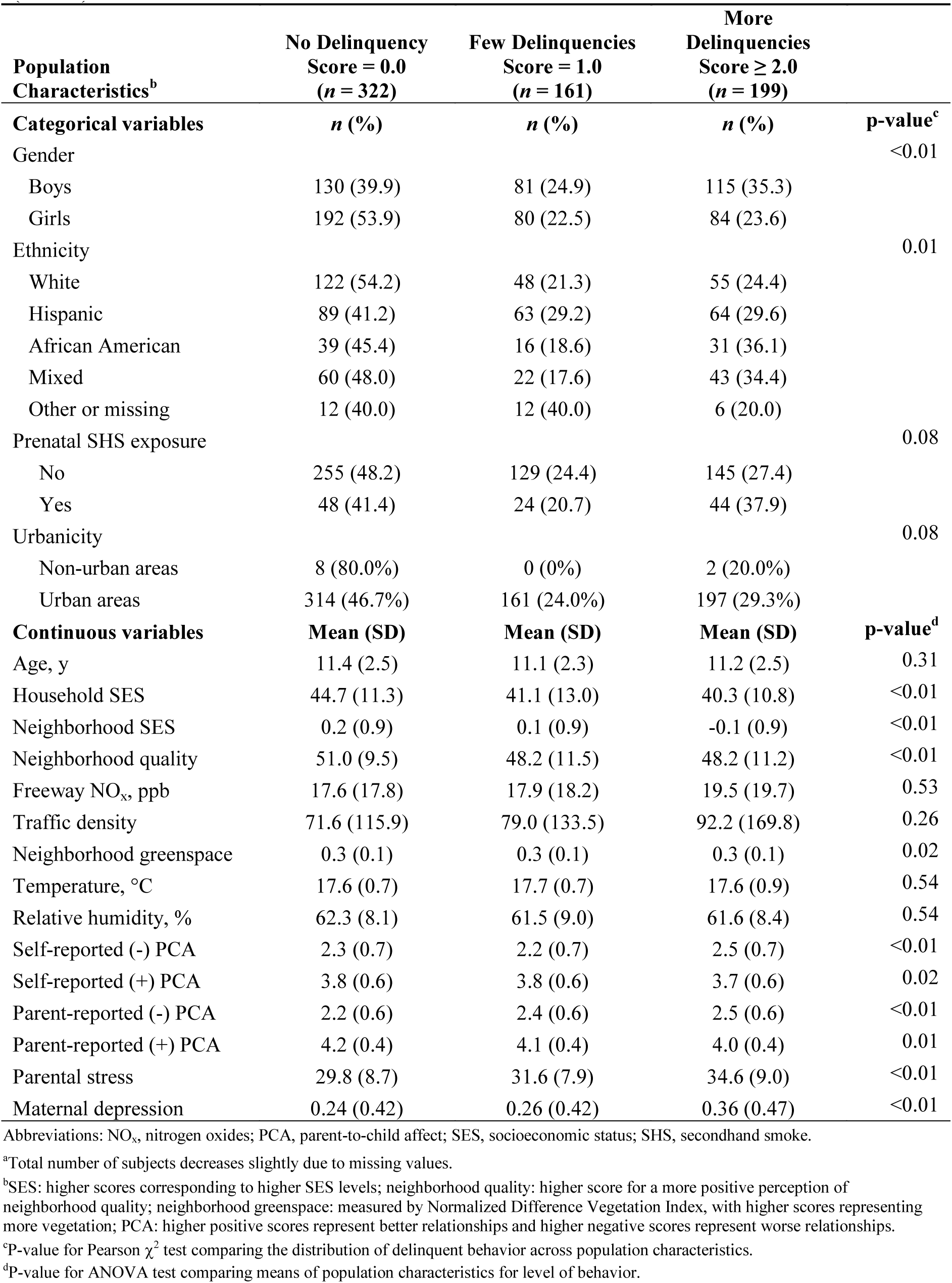
Population Characteristics in Relation to Levels of Delinquent Behavior Scores at Baseline (n=682^a^)

| **Population Characteristics<sup>b</sup>** | **No Delinquency**<br>**Score = 0.0**<br>**(n = 322)** | **Few Delinquencies**<br>**Score = 1.0**<br>**(n = 161)** | **More Delinquencies**<br>**Score ≥ 2.0**<br>**(n = 199)** |  |
| --- | --- | --- | --- | --- |
| **Categorical variables** | **n (%)** | **n (%)** | **n (%)** | **p-value<sup>c</sup>** |
| Gender |  |  |  | <0.01 |
| Boys | 130 (39.9) | 81 (24.9) | 115 (35.3) |  |
| Girls | 192 (53.9) | 80 (22.5) | 84 (23.6) |  |
| Ethnicity |  |  |  | 0.01 |
| White | 122 (54.2) | 48 (21.3) | 55 (24.4) |  |
| Hispanic | 89 (41.2) | 63 (29.2) | 64 (29.6) |  |
| African American | 39 (45.4) | 16 (18.6) | 31 (36.1) |  |
| Mixed | 60 (48.0) | 22 (17.6) | 43 (34.4) |  |
| Other or missing | 12 (40.0) | 12 (40.0) | 6 (20.0) |  |
| Prenatal SHS exposure |  |  |  | 0.08 |
| No | 255 (48.2) | 129 (24.4) | 145 (27.4) |  |
| Yes | 48 (41.4) | 24 (20.7) | 44 (37.9) |  |
| Urbanicity |  |  |  | 0.08 |
| Non-urban areas | 8 (80.0%) | 0 (0%) | 2 (20.0%) |  |
| Urban areas | 314 (46.7%) | 161 (24.0%) | 197 (29.3%) |  |
| **Continuous variables** | **Mean (SD)** | **Mean (SD)** | **Mean (SD)** | **p-value<sup>d</sup>** |
| Age, y | 11.4 (2.5) | 11.1 (2.3) | 11.2 (2.5) | 0.31 |
| Household SES | 44.7 (11.3) | 41.1 (13.0) | 40.3 (10.8) | <0.01 |
| Neighborhood SES | 0.2 (0.9) | 0.1 (0.9) | -0.1 (0.9) | <0.01 |
| Neighborhood quality | 51.0 (9.5) | 48.2 (11.5) | 48.2 (11.2) | <0.01 |
| Freeway NO<sub>x</sub>, ppb | 17.6 (17.8) | 17.9 (18.2) | 19.5 (19.7) | 0.53 |
| Traffic density | 71.6 (115.9) | 79.0 (133.5) | 92.2 (169.8) | 0.26 |
| Neighborhood greenspace | 0.3 (0.1) | 0.3 (0.1) | 0.3 (0.1) | 0.02 |
| Temperature, °C | 17.6 (0.7) | 17.7 (0.7) | 17.6 (0.9) | 0.54 |
| Relative humidity, % | 62.3 (8.1) | 61.5 (9.0) | 61.6 (8.4) | 0.54 |
| Self-reported (-) PCA | 2.3 (0.7) | 2.2 (0.7) | 2.5 (0.7) | <0.01 |
| Self-reported (+) PCA | 3.8 (0.6) | 3.8 (0.6) | 3.7 (0.6) | 0.02 |
| Parent-reported (-) PCA | 2.2 (0.6) | 2.4 (0.6) | 2.5 (0.6) | <0.01 |
| Parent-reported (+) PCA | 4.2 (0.4) | 4.1 (0.4) | 4.0 (0.4) | 0.01 |
| Parental stress | 29.8 (8.7) | 31.6 (7.9) | 34.6 (9.0) | <0.01 |
| Maternal depression | 0.24 (0.42) | 0.26 (0.42) | 0.36 (0.47) | <0.01 |

Abbreviations: NO<sub>x</sub>, nitrogen oxides; PCA, parent-to-child affect; SES, socioeconomic status; SHS, secondhand smoke.
<sup>a</sup>Total number of subjects decreases slightly due to missing values.
<sup>b</sup>SES: higher scores corresponding to higher SES levels; neighborhood quality: higher score for a more positive perception of neighborhood quality; neighborhood greenspace: measured by Normalized Difference Vegetation Index, with higher scores representing more vegetation; PCA: higher positive scores represent better relationships and higher negative scores represent worse relationships.
<sup>c</sup>P-value for Pearson $\chi^2$ test comparing the distribution of delinquent behavior across population characteristics.
<sup>d</sup>P-value for ANOVA test comparing means of population characteristics for level of behavior.

In Table 3, we present the results of the multilevel mixed-effects models, with the regression coefficient β (95% confidence interval [CI]) expressed as the difference in delinquency scores per one interquartile range (IQR) increase of PM_2.5_ (IQRs: 4.77-, 4.93-, 5.18-, and 3.12-μg/m^3^ for average 1-, 2-, 3-year and cumulative monthly average, respectively). In the base model of age-dependent trajectory accounting for within-family/individual correlations, PM_2.5_ was associated with increased delinquent behavior (all *P*s < 0.05; Table 3). In a separate set of analyses examining interaction between PM_2.5_ and age, PM_2.5_ did not affect the age-related rate of change in delinquency (Table S3). Adjustment for gender, race, household SES, nSES, and perceived neighborhood quality only modestly decreased the effect estimates and the adverse effects remained significant (Table 3). This jointly positive confounding was primarily caused by nSES characteristics, but other covariates (family SES; race/ethnicity; neighborhood quality) also had partial contributions (Tables 1 & 2). There was a consistent pattern of increased delinquency associated with PM_2.5_, and the estimates of adverse effects were equivalent to the difference in delinquency scores between subjects 3.5-4 years apart in age (Table S4).

**Table 3.** Associations^a^ Between PM_2.5_ Exposure and Repeated Measures of Delinquent Behavior

| **Models<sup>b</sup>** | **PM<sub>2.5</sub> 1-year**<br>**prior to baseline**<br>**β (95% CI)** | **PM<sub>2.5</sub> 2-years**<br>**prior to baseline**<br>**β (95% CI)** | **PM<sub>2.5</sub> 3-years**<br>**prior to baseline**<br>**β (95% CI)** | **PM<sub>2.5</sub> average**<br>**over follow-up**<br>**β (95% CI)** |
| --- | --- | --- | --- | --- |
| Base<sup>c</sup> | 0.36* (0.12, 0.60) | 0.32* (0.08, 0.56) | 0.33* (0.08, 0.58) | 0.30* (0.09, 0.52) |
| Fully Adjusted<sup>d</sup> | 0.32* (0.06, 0.59) | 0.28* (0.02, 0.54) | 0.28* (0.01, 0.56) | 0.26* (0.02, 0.51) |
| **Sensitivity Analyses** |  |  |  |  |
| I<sup>e</sup> | 0.30* (0.03, 0.57) | 0.25 (-0.02, 0.52) | 0.25 (-0.02, 0.53) | 0.25* (0.00, 0.50) |
| II<sup>f</sup> | 0.32* (0.06, 0.59) | 0.28* (0.02, 0.54) | 0.28* (0.01, 0.56) | 0.26* (0.02, 0.51) |
| III<sup>g</sup> | 0.34* (0.07, 0.62) | 0.30* (0.03, 0.56) | 0.30* (0.02, 0.58) | 0.29* (0.04, 0.54) |
| IV<sup>h</sup> | 0.33* (0.06, 0.59) | 0.28* (0.02, 0.54) | 0.29* (0.01, 0.56) | 0.25 (-0.00, 0.50) |
| V<sup>i</sup> | 0.32* (0.05, 0.59) | 0.27* (0.01, 0.54) | 0.28* (0.00, 0.56) | 0.26* (0.01, 0.51) |
| VI<sup>j</sup> | 0.30* (0.03, 0.58) | 0.25 (-0.02, 0.52) | 0.26 (-0.02, 0.54) | 0.25 (-0.00, 0.51) |
| VII<sup>k</sup> | 0.33* (0.05, 0.60) | 0.27* (0.00, 0.54) | 0.28* (0.00, 0.57) | 0.26* (0.01, 0.51) |
| VIII<sup>l</sup> | 0.33* (0.06, 0.61) | 0.28* (0.01, 0.56) | 0.28 (-0.00, 0.57) | 0.27* (0.02, 0.52) |

Abbreviations: CI, confidence interval; PM<sub>2.5</sub>, particulate matter with aerodynamic diameter < 2.5 μm.
<sup>a</sup>The results of the multilevel mixed-effects models with the regression coefficient (95% CI) representing the difference in parent-reported delinquent behavior scores across the interquartile range of PM<sub>2.5</sub> (4.77-, 4.93-, 5.18-, and 3.12-μg/m<sup>3</sup> for average 1-, 2-, 3-year and cumulative monthly average, respectively).
<sup>b</sup>n = 682 included in analyses except for sensitivity analyses VI (n=645) and VII (n=656)
<sup>c</sup>Accounting for within-family (random intercept and slope [age])/within-individual (random intercept) correlations in the model of age-dependent trajectory.
<sup>d</sup>Base + gender, ethnicity, household socioeconomic status, neighborhood socioeconomic characteristics, and neighborhood quality.
<sup>e</sup>Adjusted + freeway NO<sub>x</sub>.
<sup>f</sup>Adjusted + traffic density in 300m area.
<sup>g</sup>Adjusted + neighborhood greenspace in 250m buffer.
<sup>h</sup>Adjusted + ambient temperature.
<sup>i</sup>Adjusted + urbanicity.
<sup>j</sup>Adjusted + prenatal secondhand smoke exposure.
<sup>k</sup>Adjusted + maternal depression.
<sup>l</sup>Adjusted model using age as dichotomized variable (age ≥ 13 years vs. age < 12 years).
\*p<0.05.

Sensitivity analyses (Table 3) showed no substantial changes to the observed adverse PM_2.5_ effects for 1-year average after further accounting for freeway NO_x_, traffic densities, neighborhood greenspace, ambient temperature, urbanicity, prenatal secondhand smoke, or maternal depression. Although a few estimates for 2-year, 3-year, and cumulative monthly average became less precise and did not reach statistical significance, the results were fairly consistent as indicated by the largely overlapping confidence intervals. When dichotomized age was used in the adjusted analyses, the PM_2.5_ effects were fairly robust (Table 3: Model VIII), and interaction between PM_2.5_ and age remained non-significant (Table S2).

Delinquency increased with high levels of freeway NO_x_ in the base model (Table S5). However, adjustment of potential confounders diminished the effect estimates, which became statistically non-significant.

The associations of increased delinquency with PM_2.5_ did not vary substantially by either household SES or nSES, but were slightly stronger among boys and in families perceiving better neighborhood quality (Figure 2; Table S6). Children with lower levels of self-reported positive PCA, denoting a poorer parent-to-child relationship as compared to their counterpart, had much stronger (3-to 5-fold) adverse PM_2.5_ effects, and most of these differences were statistically significant (p < 0.05; Table S7). Interestingly, the corresponding interactions were not evident for parent-related positive PCA. The adverse PM_2.5_ effect was slightly greater in families with high (versus low) levels of negative PCA (with a worse parent-to-child relationship assessed by either children or parents), and also strengthened by parental stress or maternal depressive symptoms.

**Figure 2.**
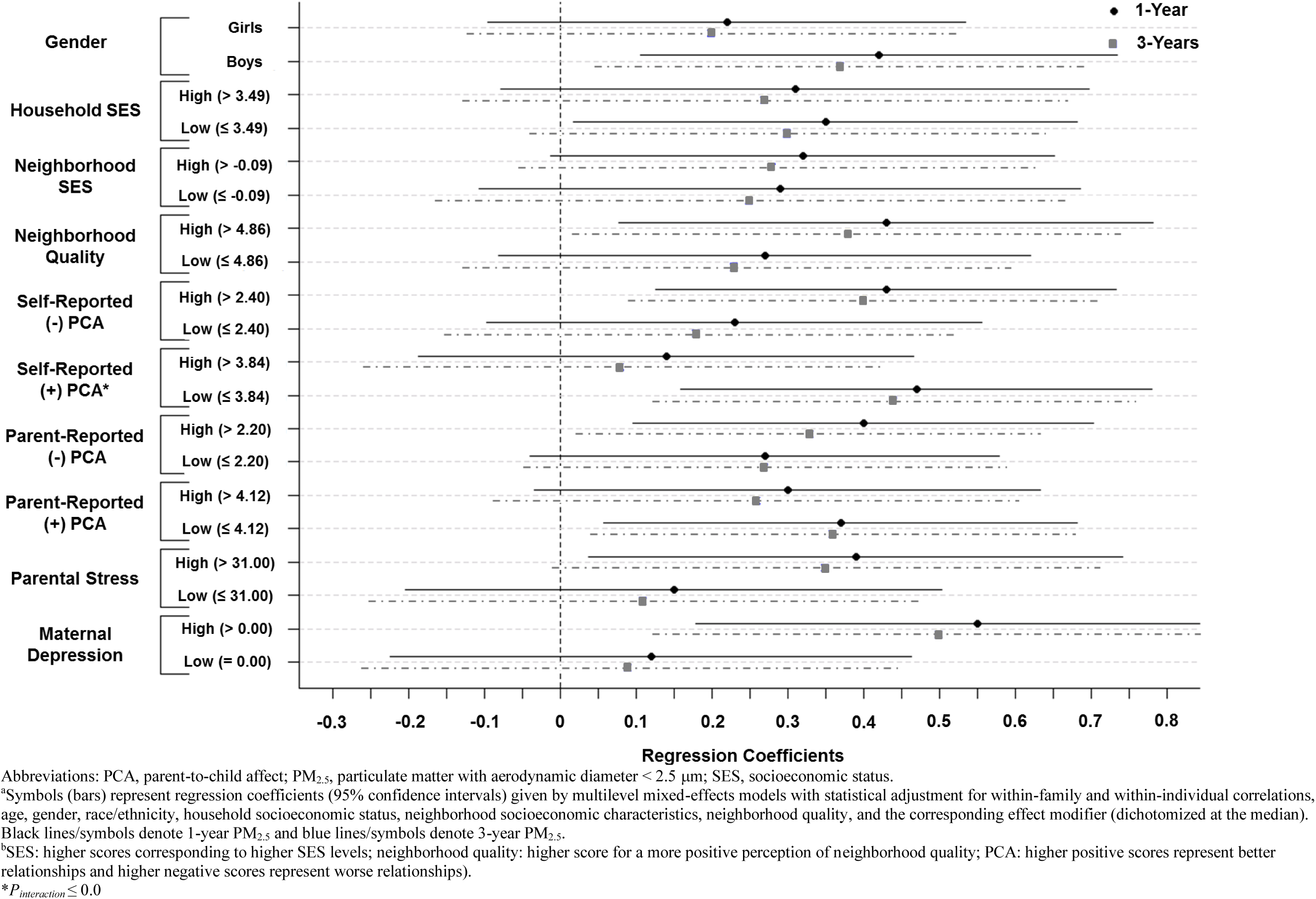
Modification of 1- and 3-year PM_2.5_ (µg/m^3^) effects^a^ on delinquent behavior by sociodemographic, neighborhood, and psychosocial factors^b^.

## Discussion

In this longitudinal study on urban-dwelling children and adolescents (ages 9-18), we found parent-reported frequency of delinquent behavior increased with residential exposure to ambient PM_2.5_ estimated at baseline (1-, 2-, and 3-year averages) and during follow-up.

The adverse PM_2.5_ effect estimates per one IQR increase were equivalent to the difference in delinquency scores between adolescents who are 3.5-4 years apart in age. These associations could not be explained by individual or household sociodemographic factors, neighborhood socioeconomic characteristics, or perceived neighborhood quality, and remained largely consistent, after statistical adjustment for freeway NO_x_, traffic density, neighborhood greenspace, meteorological factors, urbanicity, prenatal secondhand smoke, or maternal depression. The observed neurotoxic PM_2.5_ effects on delinquent behavior were strengthened by unfavorable parent-child relationships and parental psychosocial distress. Major strengths of this study include the well-characterized sample, the longitudinal study design, the extended follow-up from late childhood through adolescence, and the valid measures on psychosocial risk factors, allowing us to explore social-chemical interactions.

Our novel findings may have important public health and clinical implications. In an effort to estimate the societal impact of these results, we used the individual-level effects reported here to calculate the projected population-level effects following procedures presented in Weiss (Weiss 1988). An estimated 3.5-million adolescents reside in California (Office of Women’s Health 2009), with approximately 95% within urban areas (United States Census Bureau 2012). In our full sample, the average (±SD) delinquency score across all waves was 1.34 (± 2.06), almost identical to previous estimates for U.S. children and adolescents (Crijnen et al. 1999). A raw score at 2-SD above the mean (Achenbach and Rescorla 2001) represents a clinically significant level of CBCL-defined delinquent behavior. If the population of urban-dwelling adolescents in California (N=3.3 million) assumes the same empirical distributions (mean ± SD), 75,900 would present with delinquent behavior above the clinical range. In our study, an inter-quartile increase of PM_2.5_ was associated with a 0.26- to 0.32-point increase in delinquency scores. Given this effect size (~0.3-points), long-term PM_2.5_ exposure could presumably shift the distribution (i.e., population average) from 1.34 to 1.64. Assuming the same SD and clinically significant cut-off applies to this hypothetical scenario, this would result in an additional 16,170 cases (total cases = 92,070; a 22.4% increase) of clinically significant delinquency among California adolescents alone.

Delinquency at the clinical level at age 12-16 years is a strong predictor of adult externalizing psychopathology (Hofstra et al. 2000) and future criminal conviction (Murray and Farrington 2010). It is noteworthy that both ambient PM_2.5_ concentrations (South Coast Air Quality Management District 2013) and crime rates (Harris 2014) have been decreasing in Southern California. Therefore, it will be of great interest to conduct quasi-experimental studies (Dominici et al. 2014) investigating whether the declining PM_2.5_ due to environmental regulations might have contributed to the reduced crime rates.

Our study addresses a critical knowledge gap in the emerging area of environmental neurosciences in studying the neurobehavioral toxicity of ambient air pollutants. While several reports have linked air pollution with cognitive deficits and internalizing behavior (Block et al. 2012), the literature is scant on externalizing psychopathology. An ecological study in Ohio reported an association of adolescents’ delinquent behavior with exposure to ambient PM_2.5_ and PM_10_ (Haynes et al. 2011). In a cross-sectional study of 7 to 11 year olds in Barcelona (Forns et al. 2015), parent-reported conduct problems were associated with outdoor elemental carbon (EC; highly correlated with traffic-related PM_2.5_), but not with outdoor NO_2_. In a Cincinnati birth cohort, parent-reported conduct problems at age 7 increased with early-life EC exposure, but the adjusted association was statistically non-significant (Newman et al. 2013). Findings from these previous studies suffered from their respective methodological weaknesses (e.g., ecological design; lack of temporality in cross-sectional analyses). In our individual-level longitudinal study, residing in places with higher ambient PM_2.5_ before baseline increased delinquent behavior and such adverse effects continued during adolescence, and the observed associations remained after multiple statistical adjustments. Our study is the first to investigate whether PM_2.5_ adversely affects the trajectory of delinquent behavior in children. Although delinquent behavior became more frequent as the participants grew into adolescence, our analyses suggest no evidence for adverse PM_2.5_ effects interacting with age. If these results are substantiated in studies using similar methodological approaches, criminologists and behavioral scientists studying the effects of community and neighborhood on violent and criminal behavior may need to address the potential confounding by air pollution, as it is highly correlated with many of these factors, including poverty, neighborhood crime and violence, and neighborhood quality.

Growing experimental data have provided strong neurobiological evidence supporting adverse effects of airborne particles on increased delinquent behavior. Impulsivity and emotion dysregulation are early behavioral risk factors for externalizing psychopathology (Beauchaine 2012). Behavioral phenotypes consistent with impulsivity (preference for immediate reward; fixed interval schedule-controlled performance) were increased in mice with early-life exposures to concentrated ambient ultrafine particles (Allen et al. 2014). Such neurobehavioral effects may be mediated by particle-induced alteration of the mesocorticolimbic dopamine system (Allen et al. 2014) associated with impulsive-antisocial behavior of individuals with psychopathic personality traits. Increased depressive-like responses were also observed in mice exposed to concentrated ambient PM_2.5_ (Fonken et al. 2011) or nanoparticles from urban traffics (Davis et al. 2013). Very few human studies attempted to elucidate the neurobehavioral pathways linking air pollution exposure with externalizing psychopathology, and the reported results were mixed. In an urban Boston birth cohort, prenatal PM_2.5_ exposure was not associated with impulsivity (commission errors of the Conner’s Continuous Performance Test) at age 7 (Chiu et al. 2016). Interestingly, using the same neuropsychological tool for another urban Boston birth cohort, investigators found that traffic-related black carbon was associated with increased impulsivity ascertained at 7-14 years of age, with more prominent adverse effects in boys (Chiu et al. 2013). One cross-sectional study in Japan found that early-life exposure to suspended PM increased impulsivity-related behaviors at age 5.5 years (Yorifuji et al. 2016). In a minority urban cohort in NYC where maternal DNA adducts with polycyclic aromatic hydrocarbon (PAH) was used to measure prenatal exposure to air pollution mixture, an adverse effect on emotional self-regulation of children before age 11 was suggested, but did not adjust for secondhand smoke (Margolis et al. 2016). In a small neuroimaging subsample (n=40) selected from the same cohort, prenatal PAH exposure determined in PM_2.5_ air samples did not predict externalizing symptoms (including conduct problems) at age 7-9 (Peterson et al. 2015).

Increasing data suggest that psychological stress may enhance a child’s vulnerability to certain chemical exposures (Cooney 2011), denoting the significance of studying interactions of social-chemical stressors. In our study, the PM_2.5_-delinquency association was several-fold stronger in adolescents with psychosocial risk factors, including low-level positive PCA (self-reported), high-level negative PCA (self- and parent-reported), parental stress, and increased maternal depressive symptoms. The neurobiological mechanisms underlying the suggested psychosocial moderation of adverse neurobehavioral effects of airborne particles are unclear. Children who endure ongoing parental neglect, harsh parenting, physical or emotional abuse, poverty, or neighborhood crime and violence are in a prolonged state of toxic stress (McEwen and Tucker 2011). Growing evidence suggests chronic stress may act upon one or more of the same critical physiological pathways as air pollutants, including oxidative stress, inflammation, and autonomic disruption (Cooney 2011). One experimental model demonstrated that maternal stress during late gestation increased the offspring’s susceptibility to the deleterious effects of prenatal diesel exhaust particles exposure (Bolton et al. 2013). The researchers further posited that the synergism of air pollutants and psychosocial factors might involve their joint actions on innate immune recognition genes and downstream neuroinflammatory cascades within the developing brain. Further studies are needed to substantiate these novel hypotheses.

Our study had limitations. First, we were unable to explore early-life exposure effects because ambient PM_2.5_ data were not collected until 1999 when subjects were already 9-10 years old. Prenatal and early-childhood are critical exposure periods, which may impart much stronger neurotoxic effects, perhaps due to the higher levels of PM_2.5_ prior to 1999, so our reported association may have underestimated the true adverse PM_2.5_ effects on delinquency.

Additionally, although our analyses did not show an interaction between PM_2.5_ and age, we cannot rule out the possibility of exposure effect on early-life trajectories before age 9. Second, we used the parent-reported CBCL to assess delinquent behavior, and parents may not be aware of their children’s behaviors outside of the home. Although the Youth Self Report of CBCL was available, it is not validated for ages ≤10 and was only administered after the ages 14-15, making it unsuitable for our longitudinal analyses over ages 9-18. Third, while PM_2.5_ exposures based on spatiotemporal interpolation were well cross-validated, non-differential measurement errors in such estimates were unavoidable and likely attenuated the observed associations. Fourth, we did not study PM_2.5_ constituencies such as metals or organic species (e.g., EC). However, the correlation between EC and PM_2.5_ was very high (R^2^=0.93) in Southern California communities (Gauderman et al. 2002). Fifth, we only included subjects with complete data, potentially limiting the generalizability of our findings. However, our analyses accounted for all the major differences between our analytic sample and those excluded, and the observed associations remained in various sensitivity analyses. Sixth, we could not completely rule out the possibility of confounding by urban noise exposure. Although noise exposure has well-established adverse effects on memory functions and learning (Ferguson et al. 2013), our literature review suggests no clear evidence of a relationship between noise and externalizing behaviors (especially delinquency), and the observed PM_2.5_-effect remained statistically significant in our sensitivity analyses adjusting for traffic density. Lastly, we could not completely rule out the possibility of unmeasured or residual confounding by other environmental determinants of delinquent behavior. However, we conducted several sensitivity analyses (Table 3) and the PM_2.5_-delinquency associations persisted after statistical adjustment of multiple spatial covariates correlated with the exposure.

## Conclusions

This first longitudinal study provides supporting evidence that ambient PM_2.5_ may increase delinquent behavior before and during adolescence amongst urban-dwelling 9-18 year olds. These adverse effects may be aggravated by unfavorable parent-child relationships and parental psychosocial distress. Future studies are needed to investigate whether PM_2.5_ may affect early-life trajectories of externalizing behaviors. If the adverse neurobehavioral effects of PM_2.5_ are substantiated, the resulting knowledge may shed new light on interventions for adolescent delinquency and offer additional impetus to strengthen regulatory standards.

## Acknowledgements

This work was supported by the National Institute of Environmental Health Sciences (R21 ES022369 and F31 ES025080) and the Southern California Environmental Health Sciences Center (5P30ES007048). The USC Twin Cohort Study is funded by the National Institute of Mental Health (R01 MH058354). The authors report no conflicts of interests. The authors thank the staff and participants of the USC Twin Cohort Study for their time and effort.

## Supplementary Material

### Spatiotemporal Generalized Additive Modeling for PM_2.5_

Daily air pollution concentrations for particulate matter with aerodynamic diameter less than 2.5 μm (PM_2.5_) were acquired from the US Environmental Protection Agency’s Technology Transfer Network (Environmental Protection Agency) for the years 2000 through 2014. For exposure model development, we aggregated the daily concentrations from the monitoring sites located in our study domain to monthly averages. A generalized additive model (GAM) (Wood 2006) was fit to generate exposure estimates at subjects’ residences. The GAM is an extension of linear regression whereby spline basis functions are used to allow for smooth (non-linear) relationships between the predictor and response variables. To predict monthly PM_2.5_ at the subject locations, we incorporated smooth functions of space and time in the model

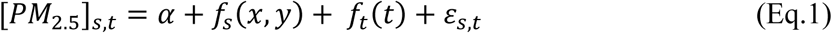

where [*PM*_2.5_]_*s,t*_ is the PM_2.5_ concentration (μg/m^3^), α is the intercept, ƒ_s_(x, y) is a 2-dimensional thin plate spline for space, s, referenced by geographic location x and y, ƒ_*t*_(t) is a cubic regression spline for time, t, referenced by calendar month, and ɛ_*s,t*_ is the iid N(0,σ^2^) residual error.

Including only those sites that met a 75% data completeness criterion, there were 25 monitoring locations in our study area with which we developed the spatiotemporal GAM exposure model. Initial model diagnostics indicated that shorter five-year GAM models generated more accurate exposure predictions than modeling the full time series in one model. Furthermore, to alleviate problems predicting on the boundaries of the time series, we split up the data into five-year segments with overlapping years (i.e. 2000-2005, 2004-2008, 2007-2011, 2010-2014) and fit separate GAM models to each segment. The average model R^2^ for the 5 five-year GAM models was 0.71. The appropriate segmented model was then used to predict PM_2.5_ concentrations at the geocoded home locations of the subjects. A monthly time-series of PM_2.5_ data from 2000-2014 was created and the monthly estimates were aggregated to represent the PM_2.5_ estimates 1-, 2-, and 3-years preceding baseline (i.e., the first valid CBCL assessment), as well as the cumulative exposure over follow-up. These different exposure estimates allow us to examine whether the putative effect might have started before and continued into the adolescence.

### CALINE Freeway NO_x_

Annual average daily traffic volumes and heavy-duty diesel truck percentage on roadways were obtained from the California Department of Transportation (Caltrans) milepost data for freeways and numbered state highways by year for 2000-2012 (CALTRANS 2010). The volumes were assigned to the ESRI Street Map Premium roadway network (www.esri.com). The extrapolated traffic count data provided 100% coverage on freeways and highways.

Estimates of the contributions of local on-road motor vehicle emissions to air quality were obtained using the CALINE4 Gaussian line-source dispersion model (Benson 1992). The CALINE4 dispersion model uses the residential locations, roadway geometry, traffic volume and emission rate by roadway link, and meteorological conditions as inputs. The model was applied to estimate annual average ambient concentration of nitrogen oxides (NO_x_) from local (within 5 km) traffic at each residence for the year of each subject’s wave test. Vehicle emission factors for each roadway segment in each year were obtained from the California Air Resources Board's EMFAC2011 model (California Air Resources Board 2013). The average vehicle speed and heavy-duty vehicle fractions of traffic volumes on specific segments, which are input to EMFAC2011, were obtained from Caltrans (CALTRANS 2008, 2010). NO_x_ was selected rather than other primary emitted species because its emission rates are most accurate. Characteristic diurnal, day-of-week, and month-of-year traffic volume variations were obtained from Caltrans (CALTRANS 2008). Hourly surface wind speed and direction data collected from meteorological stations in or near each residence during the sampling periods were used along with hourly atmospheric stability estimates and climatological estimates of morning, afternoon, and evening mixing heights in the modeling.

For this study we focused on the estimates of the near-roadway contribution of freeway traffic because it is the single most important traffic source for subjects living in urban areas such as Los Angeles. These estimated pollutant exposures should be regarded as indicators of incremental increases due to primary emissions from local vehicular traffic on top of background ambient levels. The CALINE4-estimated freeway NO_x_ only represents the effect of the incremental contribution of local traffic to a more homogeneous community background concentration of NO_x_ that included both primary and secondary pollution resulting from medium and long range transport. This indictor has been shown to explain much of the local-scale spatial variation in annual average NO_2_ and NO_x_ concentrations in southern California (Franklin et al. 2012).

### Relevant Covariate Data

Household SES was created from a composite measure of parental education levels, occupational status, family income, and marital status (Hollingshead 1979), with higher scores corresponding to higher SES levels (range: 14-66). Quality of social environments in the residential neighborhood was assessed with parent reports of 17 items regarding criminal and gang related activities, unemployment, vandalism, and substance use that occurs in the participants’ local area. This questionnaire was developed specifically for the RFAB study, resulting in a sum score (with a higher score for a more positive perception of neighborhood quality; range: 2-75) with a high level of internal consistency (average Cronbach’s Alpha across waves: α = 0.945). For household SES and quality of neighborhood social environment, missing values (11% for household SES; 3% for neighborhood quality) were imputed using the median values.

We used the socioeconomic data from 2000 US Census to objectively define the neighborhood contextual factors, as several studies have shown that the resulting neighborhood SES (nSES) characteristics are a strong predictor of neighborhood violence and crime (Kikuchi and Desmond 2010), with higher rates of crime found in disadvantaged neighborhoods. We followed the reported procedures (Dubowitz et al. 2008) to determine the nSES index that had been used in social epidemiologic studies. Briefly, an index at the census tract level was constructed using the following six variables obtained from the 2000 Census: 1) the percentage of adults 25 years old with less than a high school education; 2) the percentage of unemployed males; 3) median household income; 4) the percentage of households with income below the poverty line; 5) the percentage of households receiving public assistance; and 6) the percentage of households with children that are headed by a female. Items were summed and standardized (mean=0; SD=1), with resulting index scores greater than 0 indicating tracts above average nSES characteristics (range: −1.50–4.06).

Prenatal secondhand smoke (Tiesler and Heinrich 2014) was used as an indicator of early-life risk factors. Subjects were classified as exposed if their mothers reported either smoking during pregnancy or being exposed to secondhand smoke based on the questionnaire data collected at RFAB entry.

Maternal depression was used as an indicator of maternal risk factors and was assessed with self-reports of 6-items from the Brief Symptom Inventory (Derogatis and Melisaratos 1983) administered at baseline. Each item was coded and scored on a five-point scale, and a mean score was calculated, with higher scores indicating more symptoms of depression.

Neighborhood greenspace was estimated by Normalized Difference Vegetation Index (NDVI) (Rhew et al. 2011), derived from MODerate-resolution Imaging Spectroradiometer (MODIS) at 250-meter resolution. We obtained 16-day time-series data in 2000-2012 from the Global Agriculture Monitoring Project (NASA/GSFC et al.), and aggregated the normalized NDVI (0-1, with higher values indicating denser vegetation) in 250-m buffers surrounding residences and over various temporal scales (1-, 2-, and 3-years preceding baseline, and cumulative exposure over follow-up) (Younan et al. 2016).

Meteorological data were obtained from the California Air Resources Board (CARB) Air Quality and Meteorological Information System. Meteorological conditions recorded at the closest site were assigned to each geocoded residence to construct monthly time-series of average ambient temperature (°C) and relative humidity (%) from 1990 to 2012. Temperature and relative humidity were averaged for the periods 1-, 2-, and 3-years preceding baseline, as well as over follow-up.

Traffic density was used as a proxy for noise exposure from urban traffic. Annual average daily traffic volumes, obtained from California Department of Transportation and TeleAtlas/GDT, were assigned to the roadways and used in GIS to map traffic density within 150- and 300-m radius buffers. Annual traffic density was assigned using 2002, 2005, 2008, and 2010 roadways and average traffic volumes for waves 1-4, respectively, and baseline estimates were used.

Urbanicity was determined by spatially joining geocoded residential addresses with the 2000 and 2010 census urban area boundary files (http://www.census.gov/geo/maps-data/data/cbf/cbf_ua.html). Locations reported prior to 2006 were assigned the 2000 census urban area designation (urbanized area; urban cluster; rural area), and locations reported during or after 2006 were assigned the 2010 designation. A binary variable (urban area vs. non-urban area) was created, whereby residences located within an urban cluster or rural area were classified as non-urban.

### Measures of potential effect modifiers

Perceived parental stress was assessed with parent reports of 13 items designed to tap how unpredictable, uncontrollable, and overloaded respondents found their lives (Loeber et al. 1991). Each item was coded and scored on a five-point scale, and a continuous raw score was created by summing across items, whereby higher scores indicated more stress. This questionnaire was administered at Waves 1 and 3 with a relatively high internal consistency (average Cronbach’s Alpha across waves: α = 0.85).

Parent-to-child affect (PCA) was assessed using an adaptation of the validated Parent Feelings Questionnaire (Deater-Deckard 1996), with parallel 30-item instruments administered to both the subjects and their parents to measure both Positive (25 items) and Negative Affect (5 items). Each item was coded and scored on a five-point scale, and the means of the relevant items were calculated for each respective variable, with higher scores indicating a worse relationship between the parent and child for Negative Affect and a better relationship for Positive Affect. Caregiver and self-reports were administered at waves 1, 3, and 4 with a fairly high internal consistency for Negative Affect (average Cronbach’s Alpha across waves for caregiver reports: α = 0.67; average Cronbach’s Alpha across waves for self-reports: α = 0.66) and a relatively high internal consistency (average Cronbach’s Alpha across waves for caregiver reports: α = 0.89; average Cronbach’s Alpha across waves for self-reports: α = 0.93) for Positive Affect (Tuvblad et al. 2013).

### Three-Level Mixed Effects Modeling

Multi-level linear mixed effects models were fitted to account for clustered correlations at the temporal, individual and family levels. Under a linear growth curve modeling paradigm, random effects intercept and linear slope terms were included to account for the correlation between subjects within a family, as well as the correlation across time within a subject.

Let *Y*_*ijk*_ represent the outcome for time “k”, family “i”, and twin ‘j’

*t*_*ijk*_ represent age for family i, twin j, and time k

*t*_*ijk*_, *x*_*ij*_, *x*_*j*_ represent covariates at various levels

*U*_*ij*_, *U*_*j*_ represent random intercepts at various levels

*V*_*j*_ represents random slope at the family level

Level 1: Within person differences

Time-varying covariates: age, other spatial factors

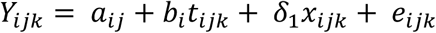

Level 2: Between person (within family) differences

Person-specific covariates: gender, neighborhood socioeconomic characteristics

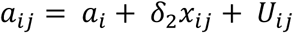

Level 3: Between family differences

Family-specific covariates: race/ethnicity, household SES, neighborhood quality, early-life and maternal risk factors

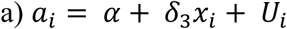

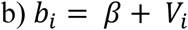

Composite linear mixed effects model:

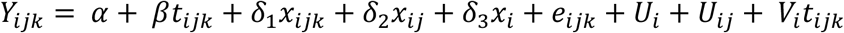

### Sensitivity and Effect Measure Modification Analyses

Based on our directed acyclic graph (DAG), age, gender, race/ethnicity, household socioeconomic status (SES), neighborhood socioeconomic (nSES) characteristics (a strong predictor of neighborhood crime rate), and self-perceived neighborhood quality were considered as cofounders in our adjusted analyses. For any observed associations that remained statistically significant in our fully adjusted models, additional analyses were conducted to evaluate whether the adjusted associations were sensitive to controlling for other spatial covariates that may be highly correlated with the exposure of interest or unmeasured early-life risk factors. Our sensitivity analyses included: 1) two-pollutant models (NO_x_ from freeway + PM_2.5_); 2) further statistical adjustment of traffic density in a 300-m buffer (proxy for neighborhood noise), neighborhood greenspace in a 250-m buffer, meteorological factors (temperature and relative humidity), prenatal secondhand smoke exposure (indicative of early-life risk factors), and maternal depression (proxy for maternal risk factors); and 3) reconstructing fully adjusted models with age specified as a dichotomized variable (age≥13 vs. age<12) to account for the possible non-linear effect of age.

Lastly, we conducted analyses to explore whether the observed exposure effects could be modified by sociodemographic factors (e.g., gender, household SES, and nSES), perceived neighborhood quality, parent-to-child affect (PCA) positive (parent- and self-report), PCA negative (parent- and self-report), perceived parental stress, and maternal depression. For better precision of the resulting stratum-specific estimates, continuous measures of potential modifiers of PM_2.5_ effects were dichotomized at the median (high vs. low). Interaction terms were included in the fully adjusted models, and Wald tests were used to assess whether effects of PM_2.5_ exposure on delinquency differed by gender (boys vs. girls), sociodemographic and neighborhood factors (high vs. low), and psychosocial stressors (high vs. low).

---

**Figure S1.**
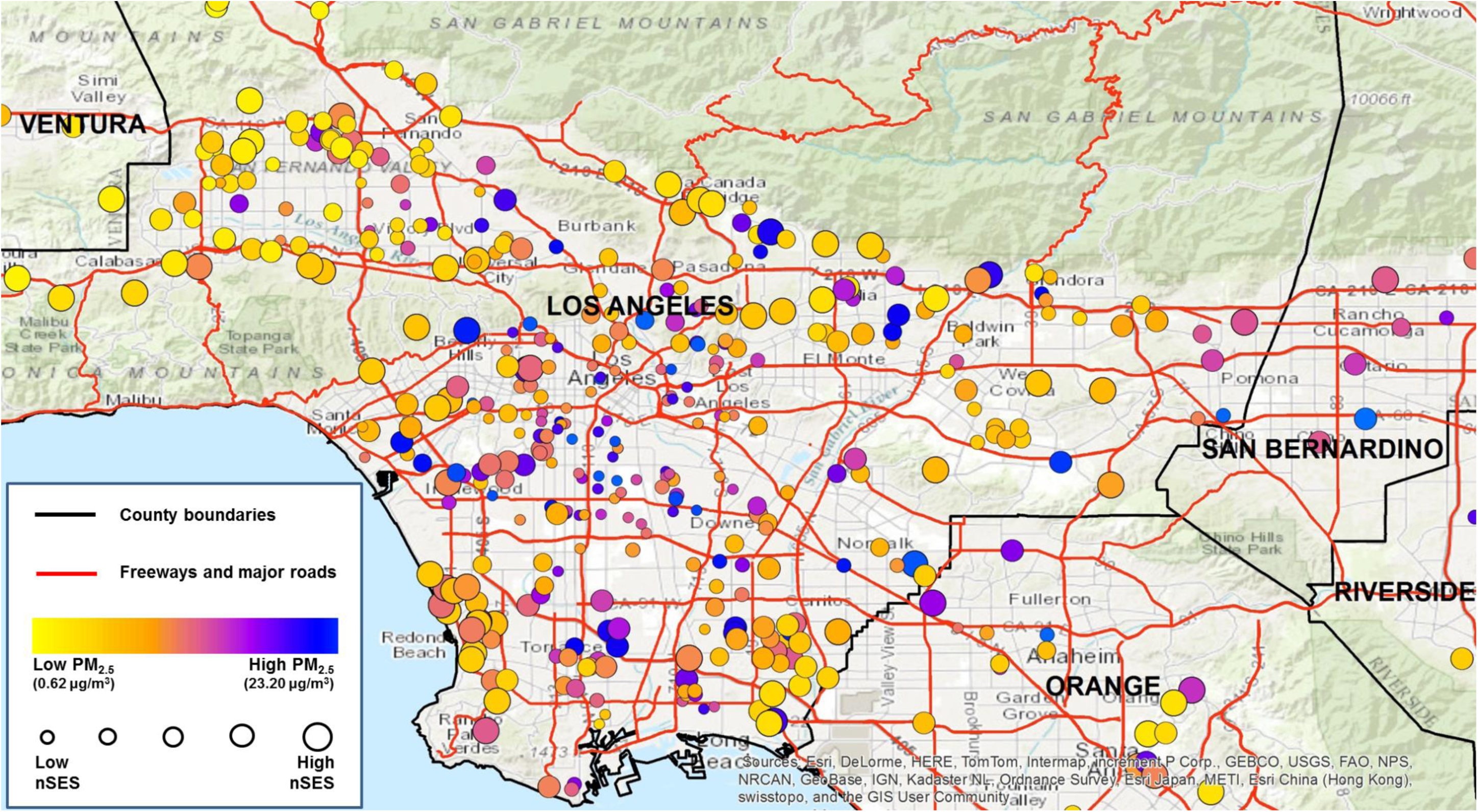
Geographic distribution of residential locations at baseline with respect to PM_2.5_ estimates and neighborhood SES levels. During the follow-up period, all families resided in Los Angeles and surrounding counties, and 68% remained at the same residential location.

**Figure S2.**
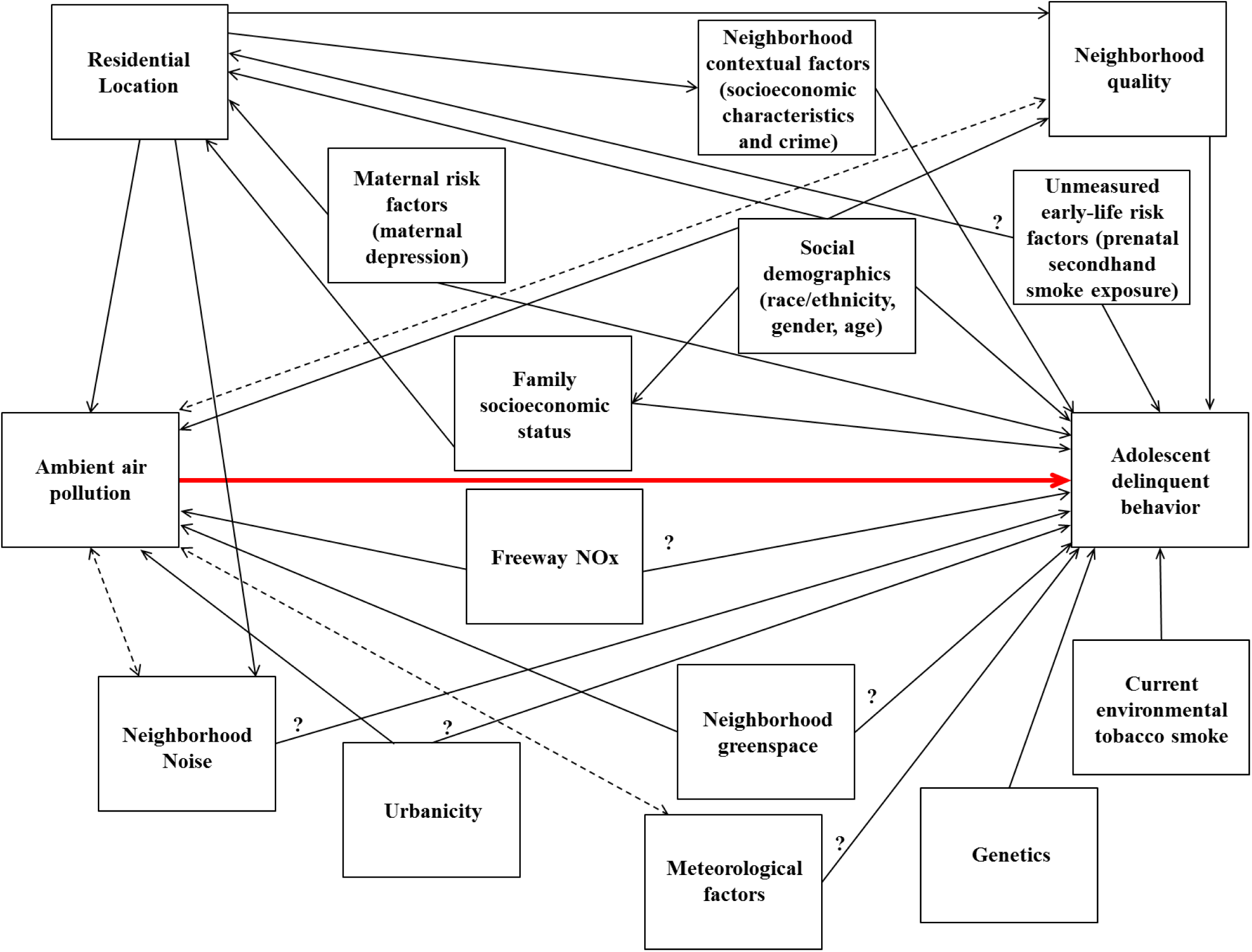
Directed acyclic graph of the relationship between ambient air pollution and delinquent behavior.

**Table S1.**
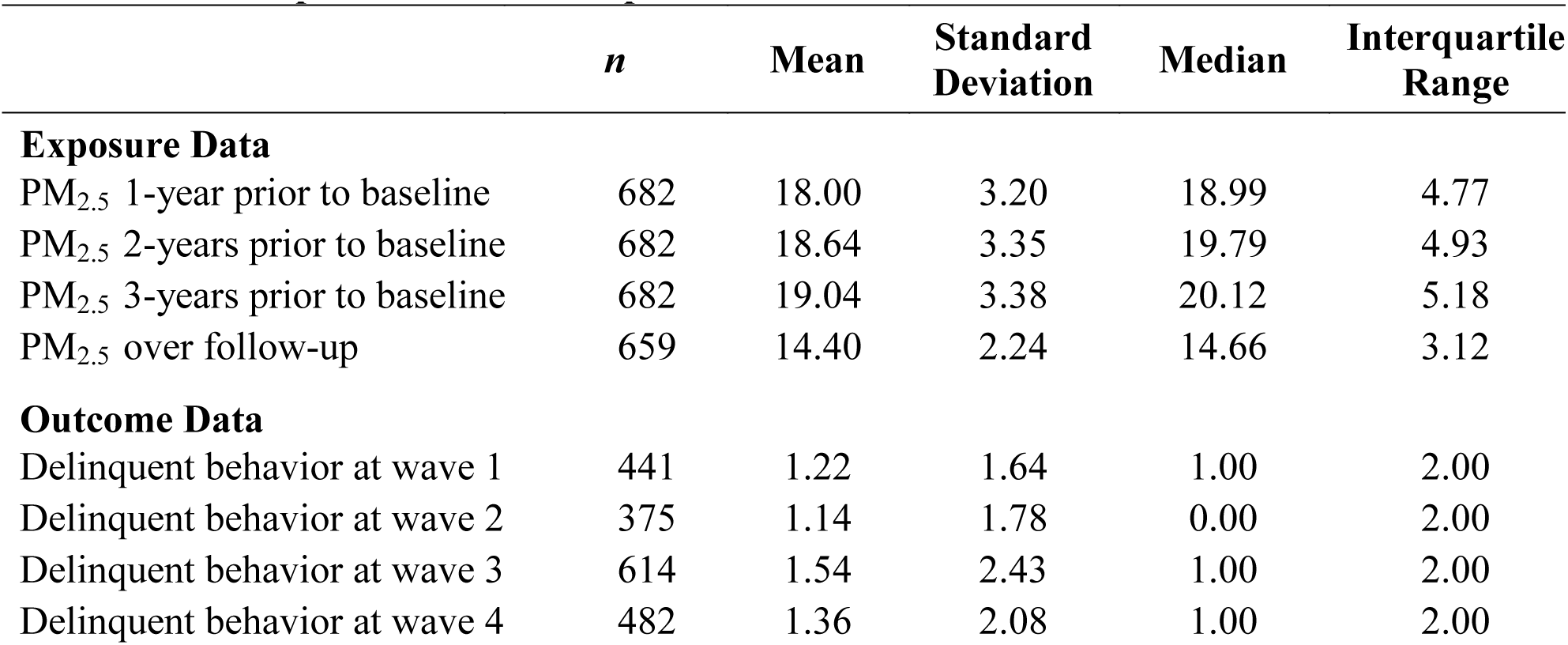
Table S1. Descriptive statistics of exposure and outcome data.

**Table S2.** Table S2. Population Characteristics of Included and Excluded Subjects

| **Population Characteristics** | **Included Subjects**<br>**(*n* = 682)** | **Excluded Subjects**<br>**(*n* = 617)** |  |
| --- | --- | --- | --- |
| **Categorical variables** | **n (%)** | **n (%)** | **p-value<sup>a</sup>** |
| Gender |  |  | 0.41 |
| Boys | 326 (51.3) | 309 (48.7) |  |
| Girls | 356 (53.6) | 308 (46.4) |  |
| Ethnicity |  |  | 0.24 |
| White | 225 (55.2) | 183 (44.9) |  |
| Hispanic | 216 (48.2) | 232 (51.8) |  |
| African American | 86 (54.1) | 73 (45.9) |  |
| Mixed | 125 (55.6) | 100 (44.4) |  |
| Other or missing | 30 (50.9) | 29 (49.2) |  |
| Baseline |  |  | <0.01 |
| Wave 1 | 441 (43.3) | 578 (56.7) |  |
| Wave 2 | 11 (84.6) | 2 (15.4) |  |
| Wave 3 | 230 (86.1) | 37 (13.9) |  |
| **Continuous variables** | **Mean (SD)** | **Mean (SD)** | **p-value<sup>b</sup>** |
| Age | 11.29 (2.42) | 9.91 (1.28) | <0.01 |
| Household SES | 42.58 (11.75) | 42.52 (11.23) | 0.93 |
| Neighborhood SES characteristics | 0.07 (0.93) | -0.04 (1.05) | 0.06 |
| Neighborhood quality | 49.43 (10.59) | 50.18 (8.87) | 0.23 |
| PM<sub>2.5</sub> 1-year prior to baseline (µg/m<sup>3</sup>) | 17.39 (3.43) | 20.76 (3.30) | <0.01 |
| Freeway NOx (ppb) | 18.22 (18.47) | 20.18 (20.21) | 0.07 |
| Delinquent behavior scores | 1.15 (1.60) | 1.26 (1.52) | 0.18 |

Abbreviations: NO<sub>x</sub>, nitrogen oxides; PM<sub>2.5</sub>, PM with aerodynamic diameter < 2.5 µm; SES, socioeconomic status.
<sup>a</sup>P-value for Pearson $\chi^2$ test comparing the distribution of subcohorts across population characteristics.
<sup>b</sup>P-value for ANOVA test comparing means of population characteristics by subcohort.

**Table S3.** Interaction Between 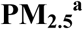 and Age on Delinquent Behavior^b^

|  | **β (95% CI)** |
| --- | --- |
| **Models for PM<sub>2.5</sub> 1-year prior to baseline (n=682)** |  |
| PM<sub>2.5</sub> 1-year prior to baseline X Age<sup>c</sup> | -0.04 (-0.29, 0.22) |
| PM<sub>2.5</sub> 1-year prior to baseline X Age<sup>d</sup> | -0.01 (-0.36, 0.33) |
| **Models for PM<sub>2.5</sub> 2-years prior to baseline (n=682)** |  |
| PM<sub>2.5</sub> 2-years prior to baseline X Age<sup>c</sup> | 0.04 (-0.21, 0.30) |
| PM<sub>2.5</sub> 2-years prior to baseline X Age<sup>d</sup> | 0.12 (-0.23, 0.48) |
| **Models for PM<sub>2.5</sub> 3-years prior to baseline (n=682)** |  |
| PM<sub>2.5</sub> 3-years prior to baseline X Age<sup>c</sup> | 0.01 (-0.22, 0.24) |
| PM<sub>2.5</sub> 3-years prior to baseline X Age<sup>d</sup> | 0.10 (-0.27, 0.47) |
| **Models for PM<sub>2.5</sub> over follow-up (n=659)** |  |
| PM<sub>2.5</sub> over follow-up X Age<sup>c</sup> | 0.01 (-0.22, 0.24) |
| PM<sub>2.5</sub> over follow-up X Age<sup>d</sup> | 0.04 (-0.26, 0.34) |

Abbreviations: CI, confidence interval; NO<sub>x</sub>, nitrogen oxides; PM<sub>2.5</sub>, PM with aerodynamic diameter < 2.5 μm; SES, socioeconomic status.
<sup>a</sup>Difference in the parent-reported frequency of child's delinquent behavior per interquartile-range increase (4.77, 4.93, 5.18, and 3.12 for 1y, 2y, 3y and cumulative average), expressed as regression coefficient (95% CI).
<sup>b</sup>Accounting for within-family (random intercept and slope [age])/within-individual (random intercept) correlations in the model of age-dependent trajectory and further adjusted for gender, ethnicity, household socioeconomic status, neighborhood socioeconomic characteristics, and neighborhood quality.
<sup>c</sup>Continuous (per 5-year increase).
<sup>d</sup>Dichotomized (age ≥ 13 years vs. age < 12 years).
\*p < 0.05.

**Table S4.** Associations Between PM_2.5_^a^ and Delinquent Behavior

|  | **Base**<br>**Model<sup>b</sup>**<br>**β (95% CI)** | **Fully Adjusted**<br>**Model<sup>c</sup>**<br>**β (95% CI)** |
| --- | --- | --- |
| **Model 1 (n=682)** |  |  |
| PM<sub>2.5</sub> 1-year prior to baseline | 0.36* (0.12, 0.60) | 0.32* (0.06, 0.59) |
| Age<sup>d</sup> | 0.09* (0.05, 0.12) | 0.08* (0.05, 0.11) |
| Girls | - | -0.49* (-0.72, -0.26) |
| **Model 2 (n=682)** |  |  |
| PM<sub>2.5</sub> 2-years prior to baseline | 0.32* (0.08, 0.56) | 0.28* (0.02, 0.54) |
| Age<sup>d</sup> | 0.08* (0.05, 0.12) | 0.08* (0.05, 0.11) |
| Girls | - | -0.49* (-0.72, -0.26) |
| **Model 3 (n=682)** |  |  |
| PM<sub>2.5</sub> 3-years prior to baseline | 0.33* (0.08, 0.58) | 0.28* (0.01, 0.56) |
| Age<sup>d</sup> | 0.08* (0.05, 0.12) | 0.08* (0.05, 0.11) |
| Girls | - | -0.49* (-0.71, -0.26) |
| **Model 4 (n=659)** |  |  |
| PM<sub>2.5</sub> over follow-up | 0.30* (0.09, 0.52) | 0.26* (0.02, 0.51) |
| Age<sup>d</sup> | 0.08* (0.05, 0.11) | 0.07* (0.04, 0.11) |
| Girls | - | -0.50* (-0.73, -0.26) |

Abbreviations: CI, confidence interval; PM<sub>2.5</sub>, PM with aerodynamic diameter < 2.5 μm.
<sup>a</sup>Difference in the parent-reported frequency of child's delinquent behavior per interquartile-range increase (4.77, 4.93, 5.18, and 3.12 for 1y, 2y, 3y and cumulative average), expressed as regression coefficient (95% CI)
<sup>b</sup>Accounting for within-family (random intercept and slope [age])/within-individual (random intercept) correlations in the model of age-dependent trajectory.
<sup>c</sup>Base + gender, ethnicity, household socioeconomic status, neighborhood socioeconomic characteristics, and neighborhood quality.
<sup>d</sup>Per 1-year increase.
\*p < 0.05.

**Table S5.** Associations Between Freeway NO_x_ and Delinquent Behavior

|  | **Base**<br>**Model<sup>b</sup>**<br>**β (95% CI)** | **Fully Adjusted**<br>**Model<sup>c</sup>**<br>**β (95% CI)** |
| --- | --- | --- |
| Freeway NO<sub>x</sub> at baseline (*n* = 682) | 0.17* (0.01, 0.33) | 0.12 (-0.04, 0.28) |
| Freeway NO<sub>x</sub> over follow-up (*n* = 659) | 0.17 (-0.00, 0.33) | 0.08 (-0.09, 0.25) |

Abbreviations: CI, confidence interval; NO<sub>x</sub>, nitrogen oxides.
<sup>a</sup>Difference in the parent-reported frequency of child's delinquent behavior per interquartile-range increase (19.15 and 15.07 for baseline and cumulative average), expressed as regression coefficient (95% CI).
<sup>b</sup>Accounting for within-family (random intercept and slope [age])/within-individual (random intercept) correlations in the model of age-dependent trajectory.
<sup>c</sup>Base + gender, ethnicity, household socioeconomic status, neighborhood socioeconomic characteristics, and neighborhood quality.
\**p* < 0.05.

**Table S6.** Modification of Adverse PM_2.5_ (µg/m^3^) Effects^a^ on Delinquent Behavior by Sociodemographics^b^

| Population Characteristics | *n* | PM<sub>2.5</sub> 1-year prior to baseline<br>β (95% CI) | PM<sub>2.5</sub> 2-years prior to baseline<br>β (95% CI) | PM<sub>2.5</sub> 3-years prior to baseline<br>β (95% CI) | PM<sub>2.5</sub> average over follow-up<sup>c</sup><br>β (95% CI) |
| --- | --- | --- | --- | --- | --- |
| **Gender** |  |  |  |  |  |
| All | 682 | 0.32* (0.06, 0.59) | 0.28* (0.02, 0.54) | 0.28* (0.01, 0.56) | 0.26* (0.02, 0.51) |
| Boys | 375 | 0.42* (0.11, 0.74) | 0.35* (0.04, 0.66) | 0.37* (0.04, 0.69) | 0.33* (0.04, 0.62) |
| Girls | 307 | 0.22 (-0.09, 0.54) | 0.21 (-0.10, 0.52) | 0.20 (-0.12, 0.53) | 0.19 (-0.10, 0.49) |
| Interaction p-value |  | p = 0.25 | p = 0.41 | p = 0.35 | p = 0.41 |
| **Household SES<sup>d,e</sup>** |  |  |  |  |  |
| All | 682 | 0.32* (0.06, 0.59) | 0.28* (0.02, 0.54) | 0.28* (0.01, 0.56) | 0.26* (0.02, 0.51) |
| High (> 3.49) | 307 | 0.31 (-0.08, 0.69) | 0.25 (-0.13, 0.63) | 0.27 (-0.13, 0.67) | 0.24 (-0.10, 0.59) |
| Low (≤ 3.49) | 375 | 0.35* (0.02, 0.68) | 0.31 (-0.01, 0.64) | 0.30 (-0.04, 0.65) | 0.27 (-0.04, 0.59) |
| Interaction p-value |  | p = 0.85 | p = 0.79 | p = 0.89 | p = 0.89 |
| **Neighborhood SES<sup>d,f</sup>** |  |  |  |  |  |
| All | 682 | 0.32* (0.06, 0.59) | 0.28* (0.02, 0.54) | 0.28* (0.01, 0.56) | 0.26* (0.02, 0.51) |
| High (> -0.09) | 350 | 0.32 (-0.02, 0.65) | 0.28 (-0.05, 0.60) | 0.28 (-0.06, 0.61) | 0.22 (-0.10, 0.53) |
| Low (≤ -0.09) | 332 | 0.29 (-0.10, 0.69) | 0.23 (-0.16, 0.63) | 0.25 (-0.17, 0.67) | 0.28 (-0.09, 0.64) |
| Interaction p-value |  | p = 0.93 | p = 0.86 | p = 0.92 | p = 0.80 |
| **Neighborhood quality<sup>d,g</sup>** |  |  |  |  |  |
| All | 682 | 0.32* (0.06, 0.59) | 0.28* (0.02, 0.54) | 0.28* (0.01, 0.56) | 0.26* (0.02, 0.51) |
| High (> 4.86) | 319 | 0.43* (0.08, 0.78) | 0.38* (0.04, 0.73) | 0.38* (0.01, 0.74) | 0.43* (0.10, 0.75) |
| Low (≤ 4.86) | 363 | 0.27 (-0.09, 0.62) | 0.21 (-0.13, 0.56) | 0.23 (-0.13, 0.59) | 0.15 (-0.17, 0.47) |
| Interaction p-value |  | p = 0.49 | p = 0.47 | p = 0.55 | p = 0.20 |

Abbreviations: CI, confidence interval; PM<sub>2.5</sub>, PM with aerodynamic diameter < 2.5 µm; SES, socioeconomic status.
<sup>a</sup>Difference in the parent-reported frequency of child's delinquent behavior per interquartile-range increase (4.77, 4.93, 5.18, and 3.12 for 1y, 2y, 3y and cumulative average), expressed as regression coefficient (95% CI).
<sup>b</sup>Accounting for within-family (random intercept and slope [age])/within-individual (random intercept) correlations in the model of age-dependent trajectory and further adjusted for gender, ethnicity, household socioeconomic status, neighborhood socioeconomic characteristics, and neighborhood quality.
<sup>c</sup>Total number of subjects decreases slightly due to missing average PM<sub>2.5</sub> over follow-up.
<sup>d</sup>Dichotomized at median.
<sup>e</sup>High group corresponds to children with high household socioeconomic status scores.
<sup>f</sup>High group corresponds to children with high neighborhood socioeconomic characteristics scores.
<sup>g</sup>High group corresponds to children of parent's with a better perception of their neighborhood.
\*p-value < 0.05 for each adjusted effect estimate.

**Table S7.** Modification of Adverse PM_2.5_ (µg/m^3^) Effects^a^ on Delinquent Behavior by Psychosocial Factors^b,c^

| Population Characteristics | *n* | PM<sub>2.5</sub> 1-year prior to baseline<br>β (95% CI) | PM<sub>2.5</sub> 2-years prior to baseline<br>β (95% CI) | PM<sub>2.5</sub> 3-years prior to baseline<br>β (95% CI) | PM<sub>2.5</sub> average over follow-up<sup>d</sup><br>β (95% CI) |
| --- | --- | --- | --- | --- | --- |
| **Self-reported (-) PCA<sup>e</sup>** |  |  |  |  |  |
| All | 655 | 0.32* (0.05, 0.59) | 0.27* (0.00, 0.54) | 0.28 (-0.00, 0.56) | 0.26* (0.01, 0.51) |
| High (> 2.40) | 330 | 0.43* (0.13, 0.74) | 0.39* (0.09, 0.69) | 0.40* (0.09, 0.72) | 0.37* (0.09, 0.64) |
| Low (≤ 2.40) | 325 | 0.23 (-0.10, 0.56) | 0.17 (-0.15, 0.50) | 0.18 (-0.15, 0.52) | 0.15 (-0.15, 0.45) |
| Interaction p-value |  | p = 0.21 | p = 0.18 | p = 0.19 | p = 0.15 |
| **Self-reported (+) PCA<sup>f</sup>** |  |  |  |  |  |
| All | 655 | 0.32* (0.05, 0.59) | 0.27* (0.00, 0.54) | 0.28 (-0.00, 0.56) | 0.26* (0.01, 0.51) |
| High (> 3.84) | 311 | 0.14 (-0.19, 0.47) | 0.08 (-0.24, 0.41) | 0.08 (-0.26, 0.42) | 0.12 (-0.18, 0.42) |
| Low (≤ 3.84) | 344 | 0.47* (0.15, 0.78) | 0.42* (0.11, 0.72) | 0.44* (0.12, 0.76) | 0.38* (0.09, 0.67) |
| Interaction p-value |  | p = 0.05 | p = 0.04 | p = 0.04 | p = 0.09 |
| **Parent-reported (-) PCA<sup>e</sup>** |  |  |  |  |  |
| All | 673 | 0.32* (0.05, 0.59) | 0.27* (0.01, 0.54) | 0.28* (0.00, 0.55) | 0.26* (0.02, 0.51) |
| High (> 2.20) | 336 | 0.40* (0.09, 0.70) | 0.35* (0.05, 0.64) | 0.33* (0.02, 0.64) | 0.32* (0.04, 0.60) |
| Low (≤ 2.20) | 337 | 0.27 (-0.04, 0.58) | 0.24 (-0.06, 0.55) | 0.27 (-0.05, 0.58) | 0.28 (-0.01, 0.56) |
| Interaction p-value |  | p = 0.46 | p = 0.52 | p = 0.70 | p = 0.78 |
| **Parent-reported (+) PCA<sup>f</sup>** |  |  |  |  |  |
| All | 673 | 0.32* (0.05, 0.59) | 0.27* (0.01, 0.54) | 0.28* (0.00, 0.55) | 0.26* (0.02, 0.51) |
| High (> 4.12) | 333 | 0.30 (-0.03, 0.64) | 0.23 (-0.10, 0.57) | 0.26 (-0.09, 0.60) | 0.20 (-0.12, 0.51) |
| Low (≤ 4.12) | 340 | 0.37* (0.06, 0.68) | 0.37* (0.06, 0.68) | 0.36* (0.04, 0.68) | 0.37* (0.08, 0.66) |
| Interaction p-value |  | p = 0.71 | p = 0.47 | p = 0.61 | p = 0.33 |
| **Perceived parental stress<sup>g</sup>** |  |  |  |  |  |
| All | 672 | 0.31* (0.05, 0.58) | 0.27* (0.01, 0.53) | 0.28* (0.00, 0.55) | 0.26* (0.01, 0.51) |
| High (> 31.00) | 347 | 0.39* (0.04, 0.75) | 0.34* (0.00, 0.69) | 0.35 (-0.01, 0.71) | 0.34* (0.02, 0.66) |
| Low (≤ 31.00) | 325 | 0.15 (-0.20, 0.51) | 0.11 (-0.24, 0.46) | 0.11 (-0.26, 0.47) | 0.11 (-0.22, 0.43) |
| Interaction p-value |  | p = 0.31 | p = 0.32 | p = 0.32 | p = 0.28 |
| **Maternal depression<sup>h</sup>** |  |  |  |  |  |
| All | 656 | 0.37* (0.09, 0.64) | 0.32* (0.05, 0.59) | 0.33* (0.04, 0.61) | 0.30* (0.05, 0.55) |
| High (> 0.00) | 327 | 0.55* (0.18, 0.92) | 0.48* (0.12, 0.84) | 0.50* (0.12, 0.88) | 0.42* (0.10, 0.74) |
| Low (= 0.00) | 329 | 0.12 (-0.22, 0.46) | 0.09 (-0.25, 0.43) | 0.09 (-0.27, 0.44) | 0.09 (-0.24, 0.42) |
| Interaction p-value |  | p = 0.07 | p = 0.09 | p = 0.09 | p = 0.12 |

Abbreviations: CI, confidence interval; PCA, parent-to-child affect; PM<sub>2.5</sub>, PM with aerodynamic diameter < 2.5 μm.<sup>a</sup>Difference in the parent-reported frequency of child's delinquent behavior per interquartile-range increase (4.77, 4.93, 5.18, and 3.12 for 1y, 2y, 3y and cumulative average), expressed as regression coefficient (95% CI).<sup>b</sup>Accounting for within-family (random intercept and slope [age])/within-individual (random intercept) correlations in the model of age-dependent trajectory and further adjusted for gender, ethnicity, household socioeconomic status, neighborhood socioeconomic characteristics, neighborhood quality, and respective covariate.<sup>c</sup>Dichotomized at median.<sup>d</sup>Total number of subjects decreases slightly due to missing average PM<sub>2.5</sub> over follow-up.<sup>e</sup>High group corresponds to worse parent-child relationship.<sup>f</sup>High group corresponds to a more favorable parent-child relationship.<sup>g</sup>High group corresponds to children of parents with more stress.<sup>h</sup>High group corresponds to children of mother's with more depressive symptoms.
\*p-value < 0.05 for each adjusted effect estimate.

## References

Achenbach, T. M., & Rescorla, L. (2001). *ASEBA school-age forms & profiles*: Aseba Burlington.

Allen, J. L., Liu, X., Weston, D., Prince, L., Oberdörster, G., Finkelstein, J. N., et al. (2014). Developmental Exposure to Concentrated Ambient Ultrafine Particulate Matter Air Pollution in Mice Results in Persistent and Sex-Dependent Behavioral Neurotoxicity and Glial Activation. Toxicological Sciences, 140(1), 160–178.

Baker, L. A., Jacobson, K. C., Raine, A., Lozano, D. I., & Bezdjian, S. (2007). Genetic and environmental bases of childhood antisocial behavior: a multi-informant twin study. J Abnorm Psychol, 116(2), 219–235, doi:10.1037/0021-843x.116.2.219.

Beauchaine, T. P. (2012). Physiological Markers of Emotional and Behavioral Dysregulation in Externalizing Psychopathology. Monographs of the Society for Research in Child Development, 77(2), 79–86, doi:10.1111/j.1540-5834.2011.00665.x.

Benson, P. E. (1992). A review of the development and application of the CALINE3 and 4 models. Atmospheric Environment. Part B. Urban Atmosphere, 26(3), 379–390.

Block, M. L., Elder, A., Auten, R. L., Bilbo, S. D., Chen, H., Chen, J.-C., et al. (2012). The Outdoor Air Pollution and Brain Health Workshop. Neurotoxicology, 33(5), 972–984, doi:10.1016/j.neuro.2012.08.014.

Bolton, J. L., Huff, N. C., Smith, S. H., Mason, S. N., Foster, W. M., Auten, R. L., et al. (2013). Maternal Stress and Effects of Prenatal Air Pollution on Offspring Mental Health Outcomes in Mice. Environ Health Perspect, 121(9), 1075–1082, doi:10.1289/ehp.1306560.

Burt, S. A. (2009). Are there meaningful etiological differences within antisocial behavior? Results of a meta-analysis. Clinical psychology review, 29(2), 163–178.

Carroll, L., Perez, M. M., & Taylor, R. M. (2014). The Evidence for Violence Prevention Across the Lifespan and Around the World: Workshop Summary: National Academies Press.

Chiu, Y. H., Bellinger, D. C., Coull, B. A., Anderson, S., Barber, R., Wright, R. O., et al. (2013). Associations between traffic-related black carbon exposure and attention in a prospective birth cohort of urban children. Environ Health Perspect, 121(7), 859–864, doi:10.1289/ehp.1205940.

Chiu, Y. H., Hsu, H. H., Coull, B. A., Bellinger, D. C., Kloog, I., Schwartz, J., et al. (2016). Prenatal particulate air pollution and neurodevelopment in urban children: Examining sensitive windows and sex-specific associations. Environ Int, 87, 56–65, doi:10.1016/j.envint.2015.11.010.

Cooney, C. M. (2011). Stress–Pollution Interactions: An Emerging Issue in Children’s Health Research. Environ Health Perspect, 119(10), a431–a435, doi:10.1289/ehp.119-a430.

Crijnen, A. A. M., Achenbach, T. M., & Verhulst, F. C. (1999). Problems Reported by Parents of Children in Multiple Cultures: The Child Behavior Checklist Syndrome Constructs. American Journal of Psychiatry, 156(4), 569–574, doi:10.1176/ajp.156.4.569.

Davis, D. A., Bortolato, M., Godar, S. C., Sander, T. K., Iwata, N., Pakbin, P., et al. (2013). Prenatal Exposure to Urban Air Nanoparticles in Mice Causes Altered Neuronal Differentiation and Depression-Like Responses. PLoS ONE, 8(5), e64128, doi:10.1371/journal.pone.0064128.

Diggle, P., Heagerty, P., Liang, K.-Y., & Zeger, S. (2002). Analysis of Longitudinal Data (2nd ed.): Oxford University Press.

Dominici, F., Greenstone, M., & Sunstein, C. R. (2014). Particulate Matter Matters. Science (New York, N.Y.), 344(6181), 257–259, doi:10.1126/science.1247348.

Ferguson, K. T., Cassells, R. C., MacAllister, J. W., & Evans, G. W. (2013). The physical environment and child development: an international review. Int J Psychol, 48(4), 437–468, doi:10.1080/00207594.2013.804190.

Fonken, L. K., Xu, X., Weil, Z. M., Chen, G., Sun, Q., Rajagopalan, S., et al. (2011). Air pollution impairs cognition, provokes depressive-like behaviors and alters hippocampal cytokine expression and morphology. Mol Psychiatry, 16(10), 987–995, 973, doi:10.1038/mp.2011.76.

Forns, J., Dadvand, P., Foraster, M., Alvarez-Pedrerol, M., Rivas, I., Lopez-Vicente, M., et al. (2015). Traffic-Related Air Pollution, Noise at School, and Behavioral Problems in Barcelona Schoolchildren: A Cross-Sectional Study. Environ Health Perspect, doi:10.1289/ehp.1409449.

Gauderman, W. J., Gilliland, G. F., Vora, H., Avol, E., Stram, D., McConnell, R., et al. (2002). Association between air pollution and lung function growth in southern California children: results from a second cohort. Am J Respir Crit Care Med, 166(1), 76–84, doi:10.1164/rccm.2111021.

Harris, K. (2014). Crime in California 2014. State of California, Department of Justice-Office of the Attorney General.

Haynes, E. N., Chen, A., Ryan, P., Succop, P., Wright, J., & Dietrich, K. N. (2011). Exposure to airborne metals and particulate matter and risk for youth adjudicated for criminal activity. Environ Res, 111(8), 1243–1248, doi:10.1016/j.envres.2011.08.008.

Hofstra, M. B., Van der Ende, J. A. N., & Verhulst, F. C. (2000). Continuity and Change of Psychopathology From Childhood Into Adulthood: A 14-Year Follow-up Study. Journal of the American Academy of Child & Adolescent Psychiatry, 39(7), 850–858, doi:http://dx.doi.org/10.1097/00004583-200007000-00013.

Howards, P. P., Schisterman, E. F., Poole, C., Kaufman, J. S., & Weinberg, C. R. (2012). “Toward a clearer definition of confounding” revisited with directed acyclic graphs. Am J Epidemiol, 176(6), 506–511, doi:10.1093/aje/kws127.

Maher, B. A., Ahmed, I. A. M., Karloukovski, V., MacLaren, D. A., Foulds, P. G., Allsop, D., et al. (2016). Magnetite pollution nanoparticles in the human brain. Proceedings of the National Academy of Sciences, 113(39), 10797–10801, doi:10.1073/pnas.1605941113.

Margolis, A. E., Herbstman, J. B., Davis, K. S., Thomas, V. K., Tang, D., Wang, Y., et al. (2016). Longitudinal effects of prenatal exposure to air pollutants on self-regulatory capacities and social competence. J Child Psychol Psychiatry, doi:10.1111/jcpp.12548.

McEwen, B. S., & Tucker, P. (2011). Critical biological pathways for chronic psychosocial stress and research opportunities to advance the consideration of stress in chemical risk assessment. Am J Public Health, 101 Suppl 1, S131–139, doi:10.2105/ajph.2011.300270.

Murray, J., & Farrington, D. P. (2010). Risk factors for conduct disorder and delinquency: key findings from longitudinal studies. Canadian Journal of Psychiatry, 55(10), 633.

Needleman, H. L., Riess, J. A., Tobin, M. J., Biesecker, G. E., & Greenhouse, J. B. (1996). Bone lead levels and delinquent behavior. JAMA, 275(5), 363–369, doi:10.1001/jama.1996.03530290033034.

Newman, N. C., Ryan, P., Lemasters, G., Levin, L., Bernstein, D., Hershey, G. K., et al. (2013). Traffic-related air pollution exposure in the first year of life and behavioral scores at 7 years of age. Environ Health Perspect, 121(6), 731–736, doi:10.1289/ehp.1205555.

Office of Women’s Health (2009). California Adolescent Health 2009. Sacramento: California Department of Health Care Services & California Department of Public Health.

Perera, F. P., Wang, S., Rauh, V., Zhou, H., Stigter, L., Camann, D., et al. (2013). Prenatal exposure to air pollution, maternal psychological distress, and child behavior. Pediatrics, 132(5), e1284–1294, doi:10.1542/peds.2012-3844.

Peterson, B. S., Rauh, V. A., Bansal, R., & et al. (2015). EFfects of prenatal exposure to air pollutants (polycyclic aromatic hydrocarbons) on the development of brain white matter, cognition, and behavior in later childhood. JAMA Psychiatry, 72(6), 531–540, doi:10.1001/jamapsychiatry.2015.57.

Petteruti, A. (2011). Education Under Arrest: The Case Against Police in Schools. Washington, D.C.: Justice Policy Institute,.

Rauh, V. A., Horton, M. K., Miller, R. L., Whyatt, R. M., & Perera, F. (2010). Neonatology and the Environment: Impact of Early Exposure to Airborne Environmental Toxicants on Infant and Child Neurodevelopment. Neoreviews, 11, 363–369, doi:10.1542/neo.11-7-e363.

Shaw, C. R., & McKay, H. D. (1942). Juvenile delinquency in urban areas. Chicago: University of Chicago Press.

South Coast Air Quality Management District (2013). Final 2012 Air Quality Management Plan. http://www.aqmd.gov/home/library/clean-air-plans/air-quality-mgt-plan/final-2012-air-quality-management-plan. Accessed April 22, 2016.

Tiesler, C. M. T., & Heinrich, J. (2014). Prenatal nicotine exposure and child behavioural problems. European Child & Adolescent Psychiatry, 23(10), 913–929, doi:10.1007/s00787-014-0615-y.

United States Census Bureau (2012). Growth in urban population outpaces rest of nation, census bureau reports. Available online: https://www.census.gov/newsroom/releases/archives/2010_census/cb12-50.html (accessed on October 28, 2015). Accessed October 28 2015.

Weiss, B. (1988). Neurobehavioral toxicity as a basis for risk assessment. Trends in Pharmacological Sciences, 9(2), 59–62, doi:http://dx.doi.org/10.1016/0165-6147(88)90118-6.

Wood, S. (2006). Generalized additive models: an introduction with R: CRC press.

Yorifuji, T., Kashima, S., Higa Diez, M., Kado, Y., & Sanada, S. (2016). Prenatal Exposure to Traffic-related Air Pollution and Child Behavioral Development Milestone Delays in Japan. Epidemiology, 27(1), 57–65.

## Reference

Benson, P. E. (1992). A review of the development and application of the CA-LINE3 and 4 models. Atmospheric Environment, 26B, 379–390.

EMFAC2011 Technical Documentation. http://www.arb.ca.gov/msei/emfac2011-documentation-final.pdf (accessed October 27, 2014) (2013).

CALTRANS (2008). 2007 California Motor Vehicle Stock, Travel and Fuel Forecast. Sacramento, CA: California Department of Transportation.

CALTRANS (2010). 2009 Annual Average Daily Traffic (AADT) for all vehicles on California State Highways. Sacramento, CA: California Department of Transportation.

Deater-Deckard, K. (1996). The Parent Feelings Questionnaire. London: Institute of Psychiatry.

Derogatis, L. R., & Melisaratos, N. (1983). The brief symptom inventory: an introductory report. Psychological medicine, 13(03), 595–605.

Dubowitz, T., Heron, M., Bird, C. E., Lurie, N., Finch, B. K., Basurto-Dávila, R., et al. (2008). Neighborhood socioeconomic status and fruit and vegetable intake among whites, blacks, and Mexican Americans in the United States. The American Journal of Clinical Nutrition, 87(6), 1883–1891.

Environmental Protection Agency, AirData. Available online: http://www.epa.gov/ttn/airs/airsaqs/detaildata/downloadaqsdata.htm (accessed July 8, 2015). Accessed July 8, 2015.

Franklin, M., Vora, H., Avol, E., McConnell, R., Lurmann, F., Liu, F., et al. (2012). Predictors of intra-community variation in air quality. J Expo Sci Environ Epidemiol, 22(2), 135–147, doi:10.1038/jes.2011.45.

Hollingshead, W. (1979). The Hollingshead four-factor index of socioeconomic status. Unpublished manuscript, Yale University, New Haven, CT.

Kikuchi, G., & Desmond, S. A. (2010). A longitudinal analysis of neighborhood crime rates using latent growth curve modeling. Sociological Perspectives, 53(1), 127–149.

Loeber, R., Stouthamer-Loeber, M., Van Kammen, W., & Farrington, D. P. (1991). Initiation, escalation and desistance in juvenile offending and their correlates. J. Crim. L. & Criminology, 82, 36.

GLAM-Global Agricultural Monitoring (2015). http://pekko.geog.umd.edu/usda/beta/data_new.php?dsRegionId=62. Accessed March 2015.

Rhew, I. C., Vander Stoep, A., Kearney, A., Smith, N. L., & Dunbar, M. D. (2011). Validation of the Normalized Difference Vegetation Index as a Measure of Neighborhood Greenness. Annals of Epidemiology, 21(12), 946–952, doi:http://dx.doi.org/10.1016/j.annepidem.2011.09.001.

Tuvblad, C., Bezdjian, S., Raine, A., & Baker, L. A. (2013). Psychopathic Personality and Negative Parent-to-Child Affect: A Longitudinal Cross-lag Twin Study. Journal of Criminal Justice, 41(5), 331–341, doi:http://dx.doi.org/10.1016/j.jcrimjus.2013.07.001.

Wood, S. (2006). Generalized additive models: an introduction with R: CRC press.

Younan, D., Tuvblad, C., Li, L., Wu, J., Lurmann, F., Franklin, M., et al. (2016). Environmental Determinants of Aggression in Adolescents: Role of Urban Neighborhood Greenspace. Journal of the American Academy of Child & Adolescent Psychiatry, 55(7), doi:http://dx.doi.org/10.1016/j.jaac.2016.05.002.

